# Cross-platform multi-laboratory screening identifies HUWE1, USP33 and USP20 as negative-regulators of autophagy or mitophagy with relevance to neurodegeneration

**DOI:** 10.64898/2026.09.07.749806

**Authors:** Maria Luisa Giudici, Emily Coode, Wing Hei Au, Clément Rey, Maria Manifava, Ovidiu-Gabriel Capatina, Gregory Aldred, Stefan Milde, Verena Zyka, Wei-Li Kuan, James Duce, Jonathan H. Clarke, Alexander J. Whitworth, Nicholas T. Ktistakis, John Skidmore

**Affiliations:** The ALBORADA Drug Discovery Institute, University of Cambridge, CB2 0AH Cambridge, UK; Signalling Programme, Babraham Institute, CB22 3AT Cambridge, UK; Medical Research Council Mitochondrial Biology Unit, University of Cambridge, CB2 0XY Cambridge, UK

**Author notes:** Joint first authors (Maria Luisa Giudici, Emily Coode). Joint senior authors (Alexander J. Whitworth, Nicholas T. Ktistakis, John Skidmore).

## Abstract

Autophagy and mitophagy are essential cellular processes implicated in neurodegenerative diseases, yet few tractable targets have been robustly validated for therapeutic modulation of these pathways. In an industry-academia consortium we carried out an extensive literature review and expert curation, leading to selection of 29 genes previously reported to enhance autophagy upon genetic or pharmacological modulation. These genes were classified based on whether downregulation or overexpression induced autophagic activity. Using siRNA knockdown, small-molecule modulators, and transient transfection approaches, we systematically screened these targets in parallel across HeLa and HEK-293 cell lines using high- and low-content phenotypic imaging. Promising candidates were further evaluated in induced pluripotent stem cell (iPSC)-derived neurons and in *Drosophila* models. Three targets emerged as top candidates: HUWE1 downregulation consistently enhanced autophagy and aggregate clearance, while USP33 downregulation promoted mitophagy. We also identified USP20, a close homologue of USP33, as an inhibitor of mitophagy, a role not previously assigned to this protein. These effects on autophagy/mitophagy were corroborated in *Drosophila*, highlighting their potential functional relevance across species. This cross-platform, multi-laboratory study establishes a framework for identifying and validating regulators of autophagy and mitophagy and provides a route which enables a comprehensive reassessment of the current target literature in this field. The validation of HUWE1, USP20, and USP33 as candidate therapeutic targets offers new opportunities for intervention in neurodegenerative diseases.

## Introduction

The pathway of autophagy is implicated in a number of human diseases, either in a tissue-specific manner or via systemic effects (Klionsky et al., 2021b; Mizushima and Komatsu, 2011). Autophagy has been proposed to function as a quality control pathway that can eliminate aggregated proteins or damaged organelles such as mitochondria and thus contribute to cellular health, particularly within the context of neurodegenerative diseases (Menzies et al., 2017; Nixon, 2013; Palmer et al., 2025). Underpinning this idea are initial observations that mice unable to activate autophagy showed ubiquitinated protein aggregates in their brains and pathology suggestive of neurodegeneration (Hara et al., 2006; Komatsu et al., 2006). In more recent work, several autophagy-regulating genes were found to be mutated in various forms of neurodegenerative disease (reviewed in (Menzies et al., 2017; Nixon and Rubinsztein, 2024). Such observations have led to the idea that molecules or treatments that enhance autophagic clearance may be of clinical benefit in various neurodegenerative diseases, especially if employed early during the pathology. Despite these observations and considerable interest in this mechanism, the translation of autophagy enhancement to the clinic has proved challenging and within the field it is recognised that the complexity of the autophagy pathway combined with the challenges of measuring the extent of autophagy in varying cellular assays has contributed to the limited progress (Choi et al., 2013; Park and Lee, 2022; Towers and Thorburn, 2016).

To tackle this challenge, we formed an industry-academia consortium consisting of three pharmaceutical companies and three academic laboratories. The consortium’s work, described in this manuscript, was undertaken to validate molecular targets whose downregulation or activation would induce autophagy in various experimental paradigms including tissue culture cells, iPSC-derived neuronal cells and *Drosophila* brains and to build understanding of the extent to which orthogonal techniques for assessing autophagic flux in cells and whole organisms generate concordant data. From an initial list of 29 genes, we identified two, HUWE1 and USP33, whose downregulation enhanced autophagy and mitochondrial autophagy (mitophagy), respectively, in all experimental settings. Subsequent experiments also revealed USP20, a homologue of USP33, as exerting a similar effect on mitophagy, a role described here for the first time. This study was significantly enabled by combining academic expertise with the drug discovery perspective of industry partners, optimizing assays for target analysis and providing more robust interrogation of these autophagy targets. In the discussion section we elaborate the benefits and challenges of this collaborative approach to target validation.

## Results

### Selection of Genes and Small Molecules for Stimulation of Autophagy and Mitophagy Pathways

A major challenge in developing clinically-viable positive modulators of autophagy lies in the limited number of well-validated, specific, and tractable molecular targets, as well as the complexity and limitations of existing disease and pathway models. While prior large-scale unbiased screens have yielded candidate regulators of autophagy (see Discussion), the goal of our consortium was to undertake a more focussed effort through a collaborative, expert-driven approach that combined academic insight and pharmaceutical discovery expertise. As a first step, we undertook a systematic, literature-informed selection process to prioritize candidate genes with reported involvement in autophagy or mitophagy. All partners proposed targets for investigation and then collectively triaged these to create a curated list of 29 targets for investigation (Table 1).

**Table 1.**
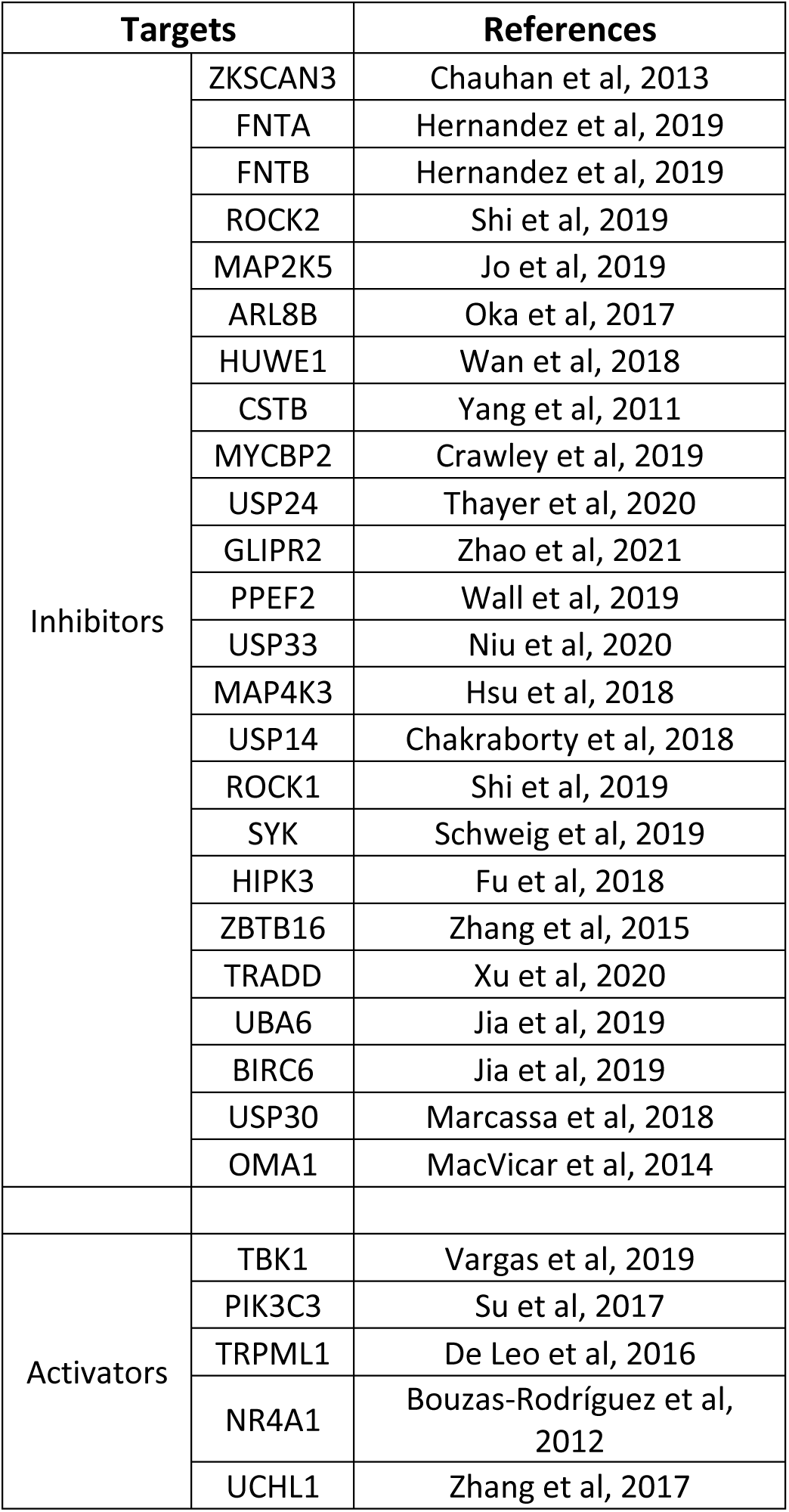
Selected targets for functional autophagy modulation.

| Targets |  | References |
| --- | --- | --- |
| Inhibitors | ZKSCAN3 | Chauhan et al, 2013 |
|  | FNTA | Hernandez et al, 2019 |
|  | FNTB | Hernandez et al, 2019 |
|  | ROCK2 | Shi et al, 2019 |
|  | MAP2K5 | Jo et al, 2019 |
|  | ARL8B | Oka et al, 2017 |
|  | HUWE1 | Wan et al, 2018 |
|  | CSTB | Yang et al, 2011 |
|  | MYCBP2 | Crawley et al, 2019 |
|  | USP24 | Thayer et al, 2020 |
|  | GLIPR2 | Zhao et al, 2021 |
|  | PPEF2 | Wall et al, 2019 |
|  | USP33 | Niu et al, 2020 |
|  | MAP4K3 | Hsu et al, 2018 |
|  | USP14 | Chakraborty et al, 2018 |
|  | ROCK1 | Shi et al, 2019 |
|  | SYK | Schweig et al, 2019 |
|  | HIPK3 | Fu et al, 2018 |
|  | ZBTB16 | Zhang et al, 2015 |
|  | TRADD | Xu et al, 2020 |
|  | UBA6 | Jia et al, 2019 |
|  | BIRC6 | Jia et al, 2019 |
|  | USP30 | Marcassa et al, 2018 |
|  | OMA1 | MacVicar et al, 2014 |
| Activators | TBK1 | Vargas et al, 2019 |
|  | PIK3C3 | Su et al, 2017 |
|  | TRPML1 | De Leo et al, 2016 |
|  | NR4A1 | Bouzas-Rodríguez et al, 2012 |
|  | UCHL1 | Zhang et al, 2017 |

In our target selection, we did not focus on proteins and complexes that constitute core components of the autophagy machinery (including autophagy receptors and autophagosome biogenesis factors) as generally these are either already being targeted or are not considered to be easily druggable. Instead, we sought potential regulators of the process either of non-selective autophagy or mitophagy. Importantly, our selection emphasized druggable targets, typically whose mechanistic roles in autophagy were reported but not yet well characterized across diverse cellular systems, particularly in neuronal models relevant to neurodegeneration (Table 1). Of note, mitophagy modulators are of particular interest in neurodegeneration (Antico et al., 2025). To identify potentially novel and tractable regulators, we also excluded canonical, extensively studied modulators such as mTORC1, VCP, PINK1, and Parkin. Target inclusion was guided by evaluation of independent studies reporting gain- or loss-of-function phenotypes, and we classified targets based on whether their reported modulation enhanced or suppressed autophagic activity. In each case, based on the team’s combined drug discovery experience, we judged that it would be feasible to develop a therapeutic that manipulated the target’s activity to increase autophagic flux. This functional annotation informed the design of downstream cellular experiments, determining whether to pursue target silencing, overexpression, or pharmacological modulation (activators or inhibitors, Table 1). This mode-of-action framework also guided the prioritization and selection of tool compounds for phenotypic validation in subsequent studies.

To identify pharmacological modulators of autophagy complementing the gene target selection, we curated a focused library of known small-molecule modulators of the targets, all of which were commercially available. This approach allowed us to assemble a set of molecules with a reasonable likelihood of producing interpretable phenotypic responses in imaging-based autophagy assays. A list of the selected compounds and their targets is provided in Table 2.

**Table 2.** Selected compounds reported to interact with the protein products of the investigated genes used for pharmacological modulation of autophagy pathways.

| Mode | Compound | Target | IC50 (nM) | References |
| --- | --- | --- | --- | --- |
| Inhibitors | Neratinib | MAP4K3 | 7.7 (Kd) | Rabindran et al, 2004 |
|  | Tipifarnib | FNTA | 0.86 | End et al, 2001 |
|  | IU1 | USP14 | 4000 | Lee et al, 2010 |
|  | Azaindole 1 | ROCK1 | 0.6 | Kast et al, 2007 |
|  | GSK269962A |  | 1.6 | Doe et al, 2007 |
|  | Chroman 1 | ROCK2 | 0.001 | Kümper et al, 2016 |
|  | Belumosudil |  | 105 | Boerma et al, 2008 |
|  | Entospletinib | SYK | 7.7 | Currie et al, 2014 |
|  | GSK143 |  | 32 | Liddle et al, 2011 |
|  | BAY61-3606 |  | 10 | Yamamoto et al, 2003 |
|  | tBID | HIPK3 | ~1,000 | Cozza et al, 2014 |
|  | SB4614852 |  | 10,000 | Patent WO02/081728 A2 |
|  | BIX02188 | MAP2K5 | 4.3 | Tatake et al, 2008 |
|  | BIX02189 |  | 1.5 | Tatake et al, 2008 |
|  | BI8626 | HUWE1 | 900 | Peter et al, 2014 |
|  | BI8622 |  | 3100 | Peter et al, 2014 |
|  | ICCB-19 | TRADD | 1120 | Xu et al, 2020 |
|  | TAK-243 | UBA6 | 7 | Hyer et al, 2018 |
|  | Competitor_USP30 | USP30 | 59 | Kluge et al, 2018 |
| Mode | Compound | Target | EC50 (nM) | References |
| Agonists | ML-SA1 | TRPML1 | 10,000 | Shen et al, 2012 |
|  | MK6-83 |  | 110 | Chen et al, 2014 |
|  | SF-22 |  | 510 | Chen et al, 2014 |
|  | SF-51 |  | 30,000 | Chen et al, 2014 |
|  | Cytosporone B | NR4A1 | 1680 (Kd) | Liu et al, 2010 |
|  | DIM-C-pPhOCH3 |  | 10,000 | Chintharlapalli et al, 2005 |
|  | Celastrol |  | 292 (Kd) | Hu et al, 2017 |

To establish a comprehensive framework for identifying robust regulators of autophagy and mitophagy, we adopted a multi-step process integrating high-throughput screening, orthogonal mechanistic validation, and cross-model replication (Fig 1 for overall description of strategy).

**Figure 1.**
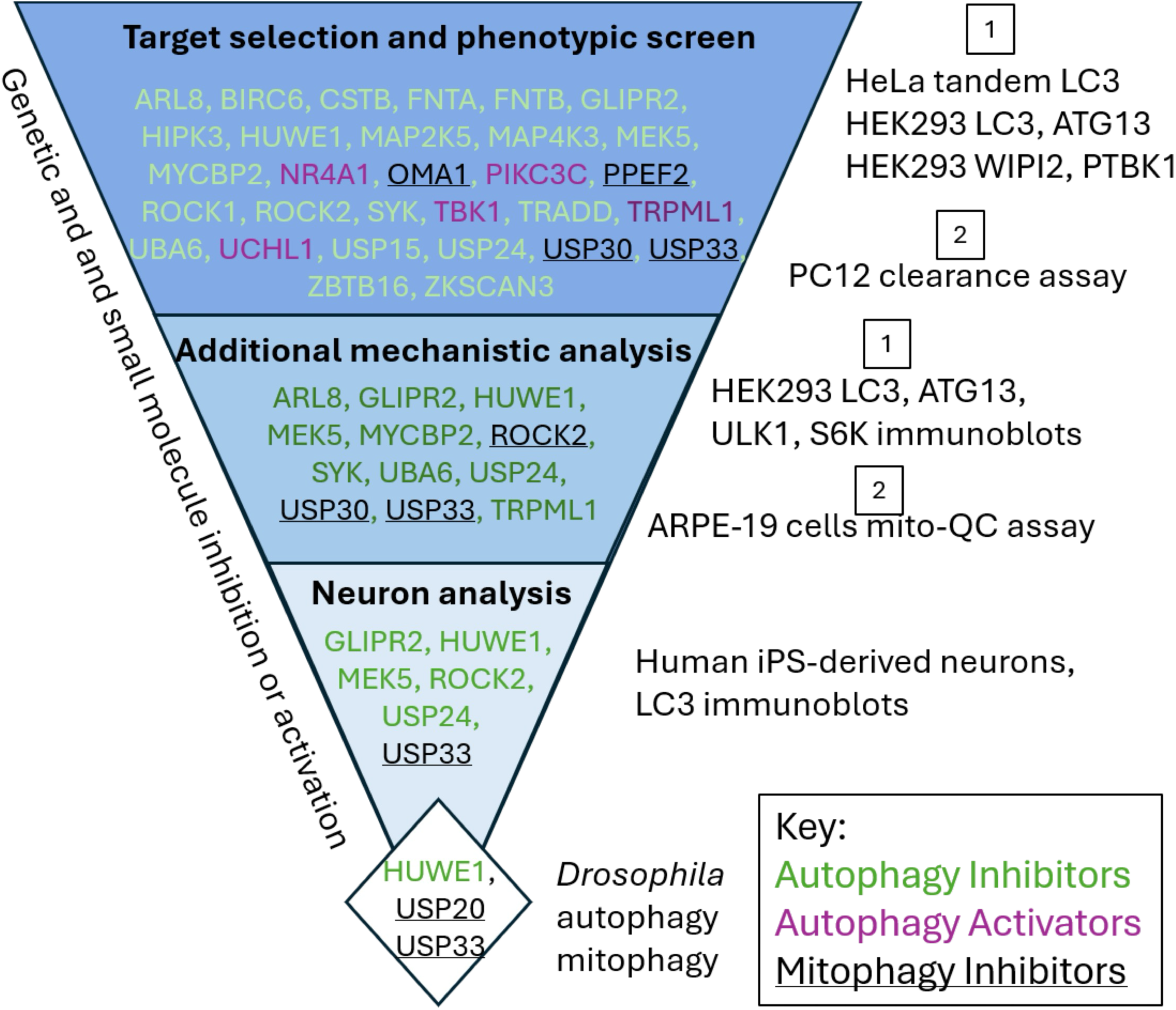
List of targets examined in this work and experimental strategy. The consortium selected 29 targets to be evaluated for their effects on autophagy. Originally all targets were examined in parallel in HeLa and HEK293 cells as indicated as well as in a protein aggregate clearance assay in PC12 cells. Targets with good responses were further analysed in more detailed biochemical assays in HEK293 cells and a subset was also analysed for their role in mitophagy in ARPE-19 cells. Positive hits from these steps were further analysed in human iPSC-derived neurons and, finally, the very top targets were analysed in *Drosophila* animals.

### Multi-platform Screening Identifies Autophagy-Modulating Targets

To investigate if these reported regulators of autophagy showed a similar phenotype in our hands, we adopted a strategy based on two different approaches that balanced scalability and mechanistic insights. First, we conducted a high-throughput, image-based screen in HeLa cells stably expressing the mRFP-GFP-LC3 reporter, a rapid and scalable assay to analyse a large number of targets but known to carry a higher level of technical noise (Fig 1). In this system, autophagosomes (positive for GFP and RFP) are distinguished from autolysosomes (positive for RFP only), enabling quantification of autophagic flux, because the GFP of the reporter is quenched in autolysosomes (Kimura et al., 2007). An automated image analysis pipeline was implemented to systematically segment cells, detect LC3 puncta, and quantify autophagosome and autolysosome populations across conditions. Cells were transfected with siRNA (SMARTpool format) targeting 24 negative regulator candidate genes. siRNA knockdown of mTOR and ATG13 served as positive and negative controls, respectively, and these performed as expected - mTOR knockdown increased autophagy, while ATG13 silencing blocked autophagosome formation - validating the assay system (Fig. 2).

**Figure 2.**
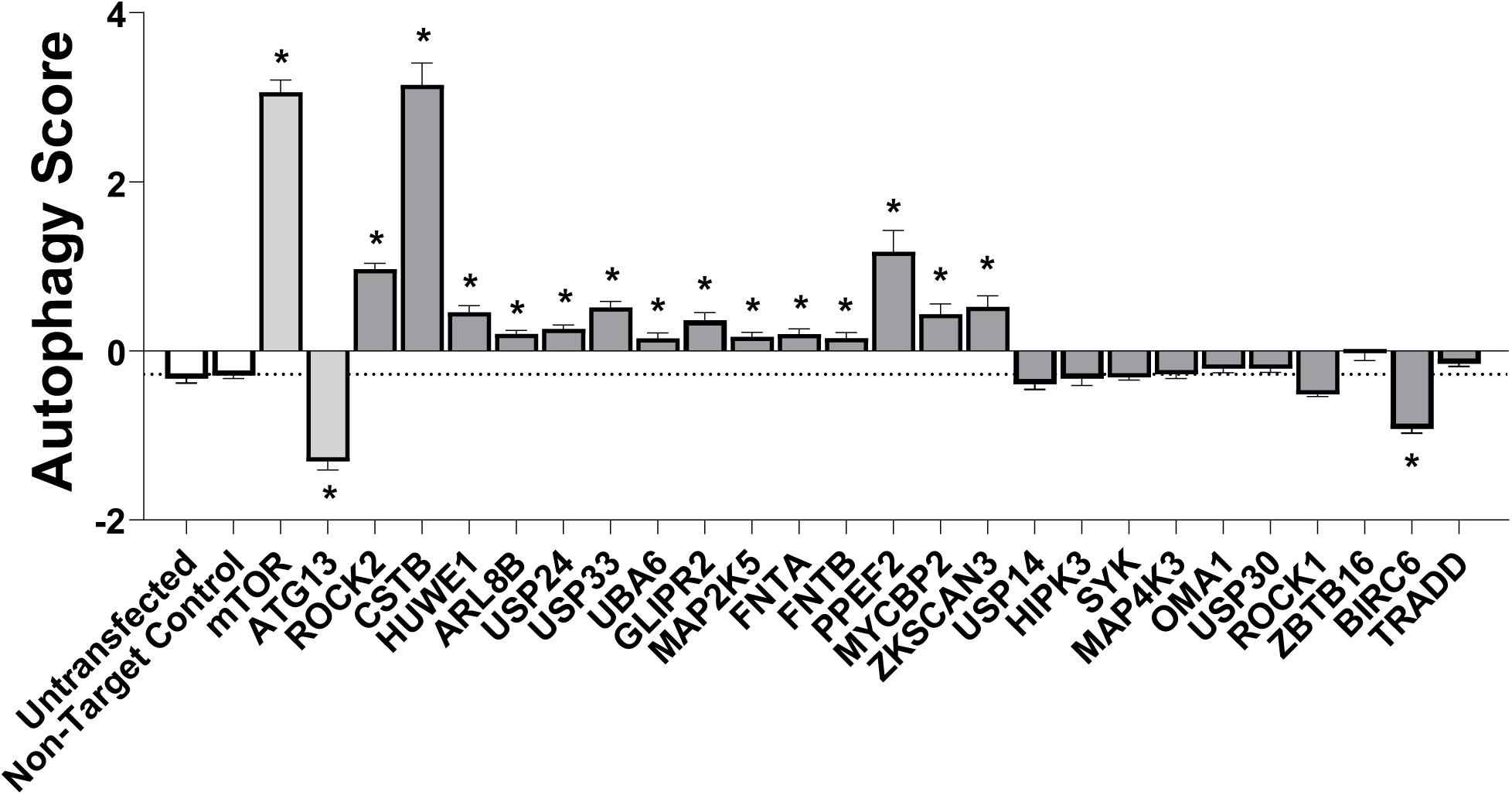
Autophagy response upon siRNA-mediated inhibition of various genes in HeLa cells and in HEK293. A. Quantification of autophagic flux in HeLa cells stably expressing the mRFP-GFP-LC3 reporter, following 96-hour 25 nM siRNA transfection. mTOR (positive control), ATG13 (negative control), non-targeting control (NTC), and non-transfected cells were included on every plate. All data are from two independent experiments, each performed across four plates, with quadruplicate wells per condition. Each plate included GFP Duplex I control siRNA to confirm transfection efficiency through reporter gene knockdown (nearly 100% expression inhibition, not shown). Statistical significance was determined using ordinary one-way ANOVA followed by Dunnett’s post hoc test, comparing each condition to the non-targeting control. p < 0.05 was considered significant (*). Values are expressed as mean ± SEM.

Assessment of cellular toxicity was performed in the same assay by quantifying Hoechst-stained nuclei (Suppl Fig. 1C). Ten gene knockdowns led to a reduction in cell count, mostly in the range of 20–40%. Interestingly, the most potent inducers of autophagy, including CSTB and the positive control mTOR, also induced the greatest levels of toxicity, suggesting a possible relationship between autophagy induction and cell viability.

In this screen, 14 out of 24 gene knockdowns induced varying degrees of autophagy activation (Fig. 2). To confirm target specificity and minimize off-target effects, all positive (leading to statistically significant increase in autophagy score) SMARTpool siRNAs were deconvoluted, and each of the four individual siRNAs per gene was tested independently. Of these, 13 of the 14 genes met stringent criteria for validation, in which at least three of the four individual siRNAs reproduced the autophagy phenotype observed with the pooled siRNA (Suppl Fig. 1A, B). UBA6 showed a partial confirmation, with only two active siRNAs, and was retained as a lower confidence hit.

High-throughput assays provide limited information about the mechanism of action of the targets and may yield false positives. Because of this, we performed parallel experiments with lower-throughput assays to probe more deeply potential mechanisms of induction and to evaluate reproducibility using alternative autophagy markers. The 24 targets described above were assayed by siRNA knock-down in HEK-293 cells stably expressing GFP-ATG13 or GFP-LC3. In the GFP-ATG13 line, cells were counterstained with WIPI2, a protein involved in the early steps of autophagosome formation (Polson et al., 2010) whereas in the GFP-LC3 line we counterstained for phospho-TBK1, an activated form of the kinase TBK1 and a marker of activated mitophagy (Heo et al., 2015). The combination of these four markers was chosen to reveal genes whose downregulation affected autophagy initiation or mitophagy. The results from this screen are shown in Table 3.

**Table 3.**
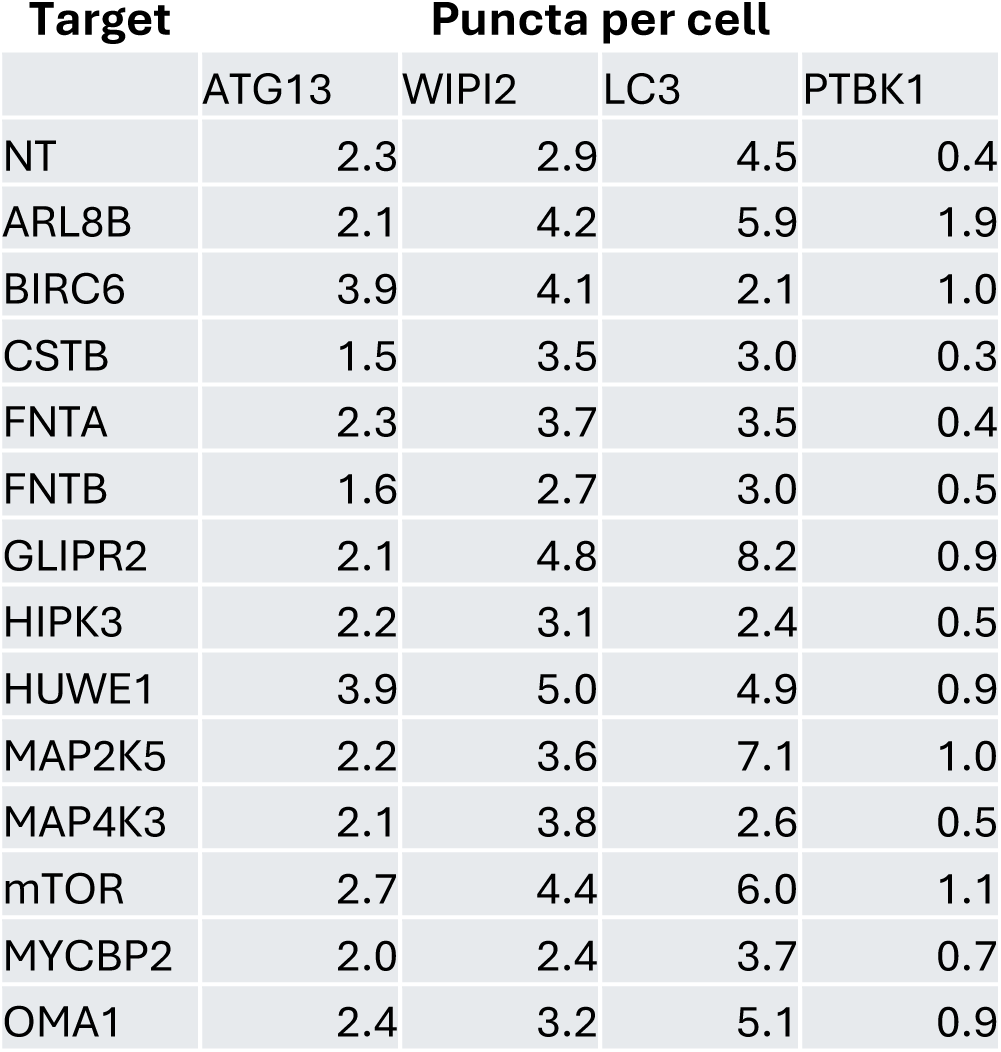
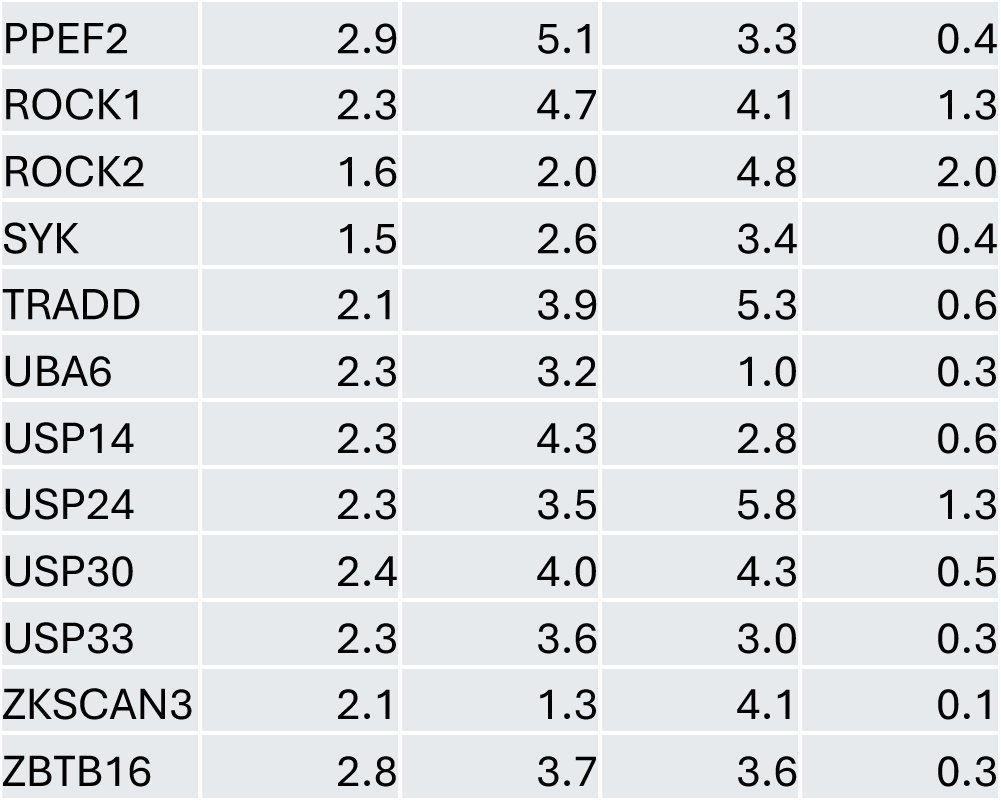
Average puncta per cell for the indicated autophagy proteins upon siRNA-mediated inhibition of various genes in HEK-293 cell lines expressing either GFP-ATG13 and counterstained with antibodies to WIPI2 or expressing GFP-LC3 and counterstained with phospho-TBK1 (Ser172). Treatment with siRNA at 100 nM was for 72 hr. NT is non-targeting control.

| Target | Puncta per cell |  |  |  |
| --- | --- | --- | --- | --- |
|  | ATG13 | WIPI2 | LC3 | PTBK1 |
| NT | 2.3 | 2.9 | 4.5 | 0.4 |
| ARL8B | 2.1 | 4.2 | 5.9 | 1.9 |
| BIRC6 | 3.9 | 4.1 | 2.1 | 1.0 |
| CSTB | 1.5 | 3.5 | 3.0 | 0.3 |
| FNTA | 2.3 | 3.7 | 3.5 | 0.4 |
| FNTB | 1.6 | 2.7 | 3.0 | 0.5 |
| GLIPR2 | 2.1 | 4.8 | 8.2 | 0.9 |
| HIPK3 | 2.2 | 3.1 | 2.4 | 0.5 |
| HUWE1 | 3.9 | 5.0 | 4.9 | 0.9 |
| MAP2K5 | 2.2 | 3.6 | 7.1 | 1.0 |
| MAP4K3 | 2.1 | 3.8 | 2.6 | 0.5 |
| mTOR | 2.7 | 4.4 | 6.0 | 1.1 |
| MYCBP2 | 2.0 | 2.4 | 3.7 | 0.7 |
| OMA1 | 2.4 | 3.2 | 5.1 | 0.9 |
| PPEF2 | 2.9 | 5.1 | 3.3 | 0.4 |
| ROCK1 | 2.3 | 4.7 | 4.1 | 1.3 |
| ROCK2 | 1.6 | 2.0 | 4.8 | 2.0 |
| SYK | 1.5 | 2.6 | 3.4 | 0.4 |
| TRADD | 2.1 | 3.9 | 5.3 | 0.6 |
| UBA6 | 2.3 | 3.2 | 1.0 | 0.3 |
| USP14 | 2.3 | 4.3 | 2.8 | 0.6 |
| USP24 | 2.3 | 3.5 | 5.8 | 1.3 |
| USP30 | 2.4 | 4.0 | 4.3 | 0.5 |
| USP33 | 2.3 | 3.6 | 3.0 | 0.3 |
| ZKSCAN3 | 2.1 | 1.3 | 4.1 | 0.1 |
| ZBTB16 | 2.8 | 3.7 | 3.6 | 0.3 |

To complement the genetic inhibition screen, we employed the same imaging approach to evaluate the effects of pharmacological modulation using the curated set of small-molecule tool compounds targeting a subset of the same candidate targets (Table 2). These compounds included inhibitors of known negative-regulators (n = 21) and agonists of potential positive regulators (n = 7), consistent with the role assigned to the target in the original publication, with the expectation that both forms of modulation would lead to increased autophagic flux.

In the HeLa mRFP-GFP-LC3 cells, each compound was tested using an eight-point concentration-response curve ranging from 6.32 nM to 20 μM. For each compound, two dose-response curves were generated: one measuring autophagy activity and the other assessing toxicity based on nuclear count (Fig 3A). This allowed us to evaluate whether autophagy was induced alongside toxicity or if there was a window between the two effects.

**Figure 3.**
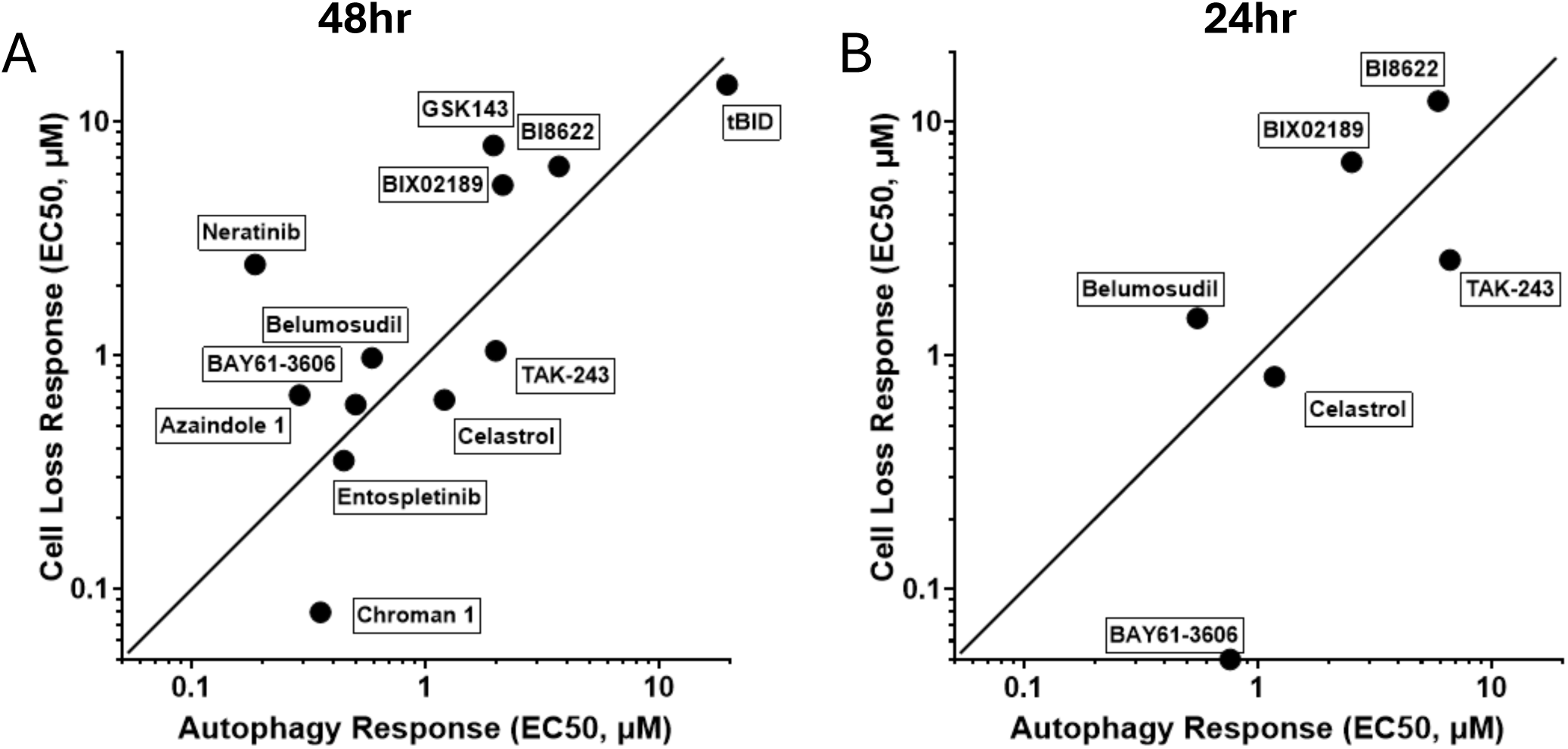
Pharmacological modulation of autophagy: evaluation of compound efficacy and toxicity. A-B. Scatter plot comparing the EC₅₀ values for autophagy induction (x-axis) and cell loss (toxicity; y-axis) for all screened compounds giving an autophagy enhancement EC₅₀. Each compound was tested in an 8-point concentration-response (6.32 nM to 20 μM) in HeLa cells expressing the mRFP-GFP-LC3 reporter following a 48-hour (A) or a 24-hour (B) exposure. Autophagy scores were normalized using DMSO (score = 0) and Torin1 (0.5 μM; score = 100) as internal controls. The line of identity (solid line) indicates equal EC₅₀ values for autophagy and toxicity. Compounds above the line exhibit effective autophagy induction at concentrations below those causing toxicity. Compounds near or below the line induce autophagy at the expense of moderate to high toxicity. The data points on the graph were collected from four separate experiments.

The correlation between autophagy activation and cell loss observed both here and in the earlier genetic screen could be due either to autophagy being activated as cell death pathways are triggered, or excessive autophagy causing cell death. We quantified the EC₅₀ values of autophagy activation versus toxicity for each compound (Fig. 3A). These EC₅₀ values were plotted against each other, and a line of identity was drawn. Compounds falling close to the identity line induced autophagy at the cost of moderate cell toxicity. Compounds positioned above the line had a lower EC₅₀ for autophagy induction than for toxicity, indicating a window between autophagy induction and cell loss. In contrast, compounds below the identity line increased autophagy only at concentrations that were also notably toxic (EC₅₀ for toxicity < EC₅₀ for autophagy) and were considered less favourable due to significant toxicity at effective doses.

From this analysis, we identified nine active compounds targeting seven negative-regulator proteins (HIPK3, HUWE1, MAP2K5, SYK, ROCK2, ROCK1, and MAP4K3) that increased autophagy without substantial toxicity. Of the seven targets identified, HUWE1, MAP2K5, and ROCK2 had also been validated in the siRNA screen, strengthening the confidence in their relevance.

To further probe compound specificity and minimize off-target effects associated with longer exposure, we repeated the compound screen using a shorter 24-hour incubation (Fig. 3B). Under these conditions, fewer compounds retained their ability to increase autophagy, however their overall toxicity profile remained largely unchanged. One exception was the SYK inhibitor BAY61-3606, which continued to induce autophagy but with increased toxicity. These results suggest that the 48-hour assay condition provides optimal sensitivity for autophagy detection, but shorter exposures may be valuable for validating compound specificity and cytotoxic thresholds.

In parallel, we employed the same two HEK-293 cell lines stably expressing GFP-ATG13 and GFP-LC3 and counterstained with antibodies to WIPI2 and phospho-TBK1 respectively to assay autophagy induction after treatment with the small molecule compounds. The results are shown in Table 4.

**Table 4.** Average number of increased puncta per cell over untreated for various autophagy proteins, for indicated pharmacological compounds (their gene targets are also shown). HEK293 cell lines expressing either GFP-ATG13 were counterstained with antibodies to WIPI2 or expressing GFP-LC3 were counterstained with phospho-TBK1 (Ser172). Cells were treated for either 1 hr or 24 hr with 10 μM of compound, shown here is the 1-hr set.

| Compound | Target | Puncta increase over untreated |  |  |  |
| --- | --- | --- | --- | --- | --- |
|  |  | LC3 | ATG13 | WIPI2 | TBK1 |
| IU1 | USP14 | 0 | 0 | 0 | 0 |
| SF-22 | TRPML1 | 2 | 0 | 0 | 0 |
| SF51 | TRPML1 | 3 | 0 | 0 | 0 |
| MK6-83 | TRPML1 | 3 | 0 | 0 | 0 |
| Cytosporone B | NR4A1 | 0 | 0 | 0 | 0 |
| ICCB-19 | TRADD | 0 | 0 | 1 | 2 |
| ML-SA1 | TRPML1 | 3 | 0 | 0 | 0 |
| Neratinib | MAP4K3 | 3 | 0 | 0 | 1 |
| Azaindole 1 | ROCK1 | 1 | 0 | 0 | 0 |
| Belumosudil | ROCK2 | 0 | 0 | 0 | 0 |
| GSK269962A | ROCK1 | 0 | 0 | 0 | 0 |
| DIM-C-pPhOCH3 | NR4A1 | 0 | 0 | 0 | 0 |
| BIX02189 | MAP2K5 | 3 | 1 | 1 | 0 |
| Tipifarnib | FNTA | 1 | 0 | 0 | 0 |
| Celastrol | NR4A1 | 0 | 0 | 0 | 0 |
| Chroman 1 | ROCK2 | 1 | 1 | 0 | 0 |
| Entospletinib | SYK | ns | ns | ns | ns |
| GSK143 | SYK | 3 | 0 | 0 | 0 |
| BIX02188 | MAP2K5 | 3 | 1 | 1 | 2 |
| TAK-243 | UBA6 | 0 | 0 | 0 | 0 |
| BAY61-3606 | SYK | 0 | 0 | 0 | 0 |
| SB4614852 | HIPK3 | 0 | 0 | 0 | 0 |
| tBID | HIPK3 | 0 | 0 | 0 | 0 |
| BI8622 | HUWE1 | 3 | 0 | 0 | 1 |
| BI8626 | HUWE1 | 2 | 1 | 0 | 0 |
| Competitor_USP30 | USP30 | 0 | 0 | 0 | 0 |
| PP242 | mTOR | 8 | 8 | 8 | 1 |

A subset of genes previously described as positive regulators of autophagy were also screened via overexpression in HeLa and HEK-293 cells using V5-tagged constructs for each gene. In HeLa cells, no hits were identified. In HEK-293 cells we observed that overexpression of TRPML1 (but no other gene) substantially increased the formation of GFP-LC3 puncta (Fig 4) but not of other markers (not shown).

**Figure 4.**
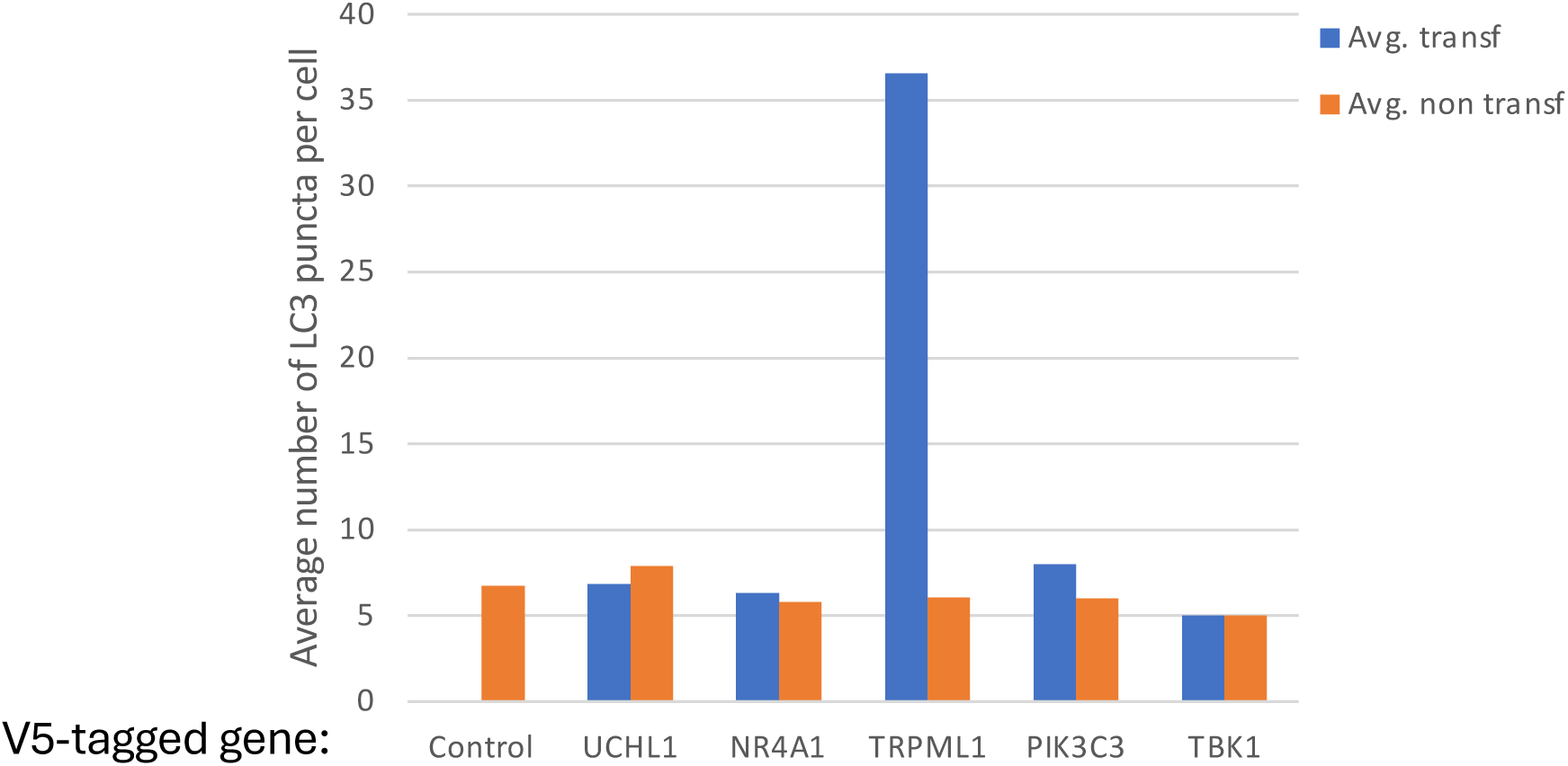
Average number of LC3 puncta per cell upon overexpression of various genes. The number of LC3 puncta in transfected (+ve) and non-transfected (-ve) HEK-293 cells were counted. Transfection was for 48 hr.

Taken together, these multi-platform screening efforts provide a comprehensive and cross-validated framework for prioritizing targets. The high-throughput siRNA screen identified 13 reproducible genes, establishing the primary dataset for downstream evaluation. Because siRNA knockdown offers high target specificity, these genetic hits served as the foundation for subsequent validation. A complementary compound screen was performed on available tool compounds corresponding to a subset of these genes; among the four genes with suitable modulators, three (ROCK2, MAP2K5, and HUWE1) reproduced the siRNA phenotype, increasing autophagic flux without excessive toxicity. These convergent results from genetic and pharmacological perturbation increased confidence in these three targets.

Of the 13 targets validated as positive in the high-throughput siRNA screen, the low-throughput siRNA screen indicated that five (HUWE1, MAP2K5, USP24, ARL8B, and GLIPR2) also increased autophagy markers in these assays, demonstrating reproducibility across models and independent markers. Of the four genes with tools compounds, ROCK2 and MAP2K5 again showed positive outcomes, with MAP2K5 consistently reproducing its phenotype under both genetic and pharmacological modulation. Overexpression studies highlighted TRPML1 as a potential positive regulator of autophagy.

Based on the reproducibility of autophagy induction across multiple cell lines and assay formats, the consortium nominated the targets HUWE1, MAP2K5, USP24, ARL8B, GLIPR2, ROCK2, and TRPML1 for further investigation which will be reported later.

### Phenotypic Image-Based High-Throughput Screening: Identification of Targets Involved in Protein Clearance

Following the identification of candidate autophagy regulators, we next assessed whether modulating these targets could influence the clearance of protein aggregates, a process highly relevant for neurodegenerative disease (Menzies et al., 2017). Target activity was altered either genetically or pharmacologically, and the resulting effects on protein clearance were evaluated using two independent, image-based assays (Suppl Fig 2).

The primary assay employed a PC12 cell line stably expressing GFP-tagged exon 1 of mutant huntingtin (Htt) containing a pathological 74-glutamine (Q74) expansion (Wyttenbach et al., 2001). Expression of the fusion protein was induced via a doxycycline-regulated promoter. Upon induction, cells accumulate both a diffuse GFP signal, reflecting soluble Htt, and discrete, brightly fluorescent inclusions corresponding to aggregated protein. Protein clearance was quantified by two parameters: (i) the average GFP intensity per cell and (ii) the proportion of cells containing one or more visible aggregates. These two readouts were integrated into a composite “clearance score”, which served as a functional readout of the target’s impact on proteostasis.

This clearance assay employed both genetic (Suppl Fig. 2A) and pharmacological (Suppl Fig. 2B) manipulation. For the genetic screen, the PC12 cells were treated with SMARTpool siRNAs. Controls included uninduced cells (baseline clearance; 100%) and induced cells transfected with non-targeting siRNA (aggregated condition; 0%). Moreover, siRNA against mTOR and ATG13 served as positive and negative controls, respectively. As expected, mTOR inhibition led to an improvement in clearance score (by about 20%), confirming that increased autophagy correlates with enhanced removal of mutant Htt aggregates. Among the candidate genes tested, siRNA-mediated knockdown of four targets (HUWE1, MYCBP2, USP30, and ZBTB16) produced modest but statistically significant increases in clearance, each in the range of ∼10%. These small but consistent effects suggest a potential, albeit limited, role in aggregate handling.

In the compound screen, each tool compound was tested using a full 8-point concentration-response format, with a 48-hour incubation in the continued presence of doxycycline. Clearance scores were scaled relative to uninduced cells (score = 100) and DMSO-treated, induced cells (score = 0). For each compound, a pair of curves was generated: one representing the clearance score, and the other measuring cell number to assess compound toxicity. As in the autophagy compound screen, EC₅₀ values were calculated for both clearance and toxicity. These values were then compared to assess whether clearance improvements occurred independently of cell death. Compounds showing clearance effects at concentrations below those causing substantial toxicity were considered more likely to act via specific modulation of proteostasis pathways.

This analysis identified four compounds that improved aggregate clearance (Suppl Fig 2B): inhibitors of SYK and ROCK1/ROCK2 (BAY61-3606 and Azaindole 1, respectively), and activators of TRAD and NR4A1 (ICCB-19 and Celastrol, respectively). It is of note that the compound Azaindole 1 is a dual inhibitor of ROCK1 and ROCK2 (both isoforms of the Rho-associated protein kinase). These effects occurred in the context of low to moderate toxicity, suggesting that some compounds may influence proteostasis through mechanisms that do not fully recapitulate genetic knockdown and highlight the utility of complementary screening approaches to uncover functionally relevant regulators of aggregate clearance.

To complement the protein clearance findings from the PC12 model, we employed a second phenotypic assay using HeLa cells transiently transfected with a 0N4R-Tau-HiBiT(P301S) construct (Suppl Fig 2C). This tau isoform, prone to misfolding and aggregation into neurofibrillary tangles(Tuck et al., 2022), was tagged with the HiBiT peptide, allowing quantification of total tau levels through luminescence upon addition of its complementing protein LgBiT in cell lysates. As with the PC12 assay, compound toxicity was assessed in parallel to discriminate genuine tau clearance effects from those secondary to cytotoxicity. EC₅₀ values were calculated for both tau clearance and cell viability and plotted relative to each other (Suppl Fig. 2C). We focused analysis on compounds with clearance EC₅₀ values below 20 μM to enrich for potent and selective modulators. Following this stringent selection, five putative negative autophagy regulators (ROCK1, MAP4K3, ROCK2, MAP2K5, and SYK) and one positive regulator (NR4A1) emerged. These findings partially overlapped with those from the PC12-based assay, which had also previously identified targets ROCK1/ROCK2, SYK, and NR4A1, highlighting both shared and distinct regulatory effects across models.

It is important to note that the PC12 assay separately quantified both soluble and aggregated forms of GFP-Htt(Q74), whereas the HiBiT-based tau assay measured total protein expression without discriminating for aggregation state. Together, these complementary approaches reinforce the identification of key modulators influencing proteostasis, offering insights into target-specific effects on protein accumulation relevant to neurodegenerative disease pathology.

Taken together, the outcomes of these siRNA and compound screens indicate the targets that most robustly influence aggregate clearance. Although four genes (HUWE1, MYCBP2, ZBTB16, and USP30) modestly improved clearance following siRNA knockdown, only HUWE1 and MYCBP2 also scored positively in the autophagy assays, but MYCBP2 displayed inconsistent activity across mitophagy readouts (see later section), diminishing its confidence level as a bona fide regulator. In contrast, the compound screen identified seven targets (ROCK2, MAP2K5, ROCK1, SYK, TRADD, and NR4A1) whose pharmacological modulation enhanced clearance, yet only ROCK2 and MAP2K5 showed supporting evidence from autophagy assays. Even so, their genetic perturbation did not phenocopy the compound effects, suggesting off-target compound activity. When considering hit robustness across both genetic and pharmacological perturbation, as well as alignment with autophagy-enhancing phenotypes, HUWE1 emerges as the strongest and most consistent regulator of aggregate clearance among the seven targets previously nominated as potential autophagy regulators.

### Mechanistic Characterisation of Candidate Regulators Identified Through Multiplatform Autophagy and Clearance Screening

To gain a deeper mechanistic understanding of the targets prioritised from the preceding high-throughput and low-throughput autophagy and clearance assays, we carried out additional functional investigations in HEK-293 cells on SYK, MAP2K5, ARL8B, TRPML1, USP24, GLIPR2, ROCK2, and HUWE1. This approach was designed to validate the role of candidate targets in autophagy within an alternative cellular context, and to leverage a series of low-throughput, mechanistically informative assays capable of distinguishing between bulk autophagy, selective autophagy (including mitophagy), and non-canonical autophagic pathways.

#### SYK

Inhibition of SYK has been reported to enhance autophagy which may provide therapeutic benefit in aggregate clearance and neurodegeneration (Krisenko et al., 2015; Schweig et al., 2019). Our data using the SYK inhibitor GSK143 had shown an increase in LC3 autophagic puncta in both HEK-293 and HeLa cells by immunofluorescence (Fig 3A and Table 4). This was confirmed in biochemical assays as an increase in autophagic LC3-II levels (Suppl Fig 3A, compare lanes in control vs GSK143 treated samples). However, addition of bafilomycin A1 did not increase levels of LC3-II further (Suppl Fig 3A, compare Baf lanes) which is generally interpreted to mean that the SYK inhibitor did not induce autophagy but blocked autophagic flux (Klionsky et al., 2021a). This was confirmed by immunofluorescence staining of endogenous LC3: although GSK143 increased LC3 puncta indicating formation of autophagic structures this did not increase further upon bafilomycin A1 treatment, indicating that GSK143 is likely to affect the clearance of autophagosomes and not their formation (Suppl Fig 3B). Furthermore, in more detailed biochemical experiments using siRNA- mediated downregulation of SYK, we did not obtain evidence that LC3-II levels increased upon significant reduction of the protein (Suppl Fig 3C). We concluded from the above that the LC3 signal observed with the GSK143 inhibitor likely represents an off-target effect which in any case affects autophagosome clearance and not formation.

#### MAP2K5

Previous work has suggested that inhibition of MAP2K5 induces autophagy, likely via an mTOR-independent mechanism (Jo et al., 2019). We also observed that treatment of cells with two related MAP2K5 inhibitors (BIX02188 and BIX02189) resulted in significant induction of autophagy as judged by increased puncta formation of two early autophagy proteins FIP200 and WIPI2 (Suppl Fig 4A, B). However, because these two compounds are structurally similar (and the data could therefore not be considered entirely independent) we tested a third inhibitor, GW284543, which did not recapitulate the results (data not shown). In addition, we examined mTOR activation in response to BIX02188 and BIX02189 and found that these compounds inhibit mTOR-mediated phosphorylation of S6 kinase and of ULK1 (Suppl Fig 4C). Combined, these data suggest that MAP2K5 is a less robust target for autophagy initiation, and that the strong effects seen with BIX02188 and BIX02189 are likely off-target and affect mTOR activation state, a known regulator of autophagy. To further investigate this point, we carried MAP2K5 into the next phase of analysis in iPSC-derived neuronal cells as a low-confidence target (see below).

#### ARL8B

Previous work suggested a role for the small GTPase ARL8B in lysosome positioning and cargo delivery (Korolchuk et al., 2011). These functions were shown to indirectly affect autophagy at the level of lysosome-autophagosome fusion, notably in neuronal cells (Adnan et al., 2020). In our assays, downregulation of ARL8B showed increased formation of autophagic structures positive for early autophagy proteins especially WIPI2 (Table 3). In subsequent experiments assaying formation of LC3-II by immunoblots we showed that cells downregulated for ARL8B had a mild increase in basal autophagy and in autophagy following mTOR inactivation with PP242, but this was not accompanied by a significant enhancement of the response when bafilomycin A1 was used to block autophagosome-lysosome fusion (Suppl Fig 5). We concluded from these data that the effects of ARL8B in the autophagy pathway are complex and likely involve both the induction and the maturation steps.

#### TRPML1

The calcium permeable ion channel TRPML1 that is located on late endosomes and lysosomes has been implicated in autophagy, both at the initiation as well as the maturation step (Scotto Rosato et al., 2019). Using a variety of agonists, we verified that activation of the channel produces a strong LC3 response which however was not accompanied by a response from early autophagy proteins such as ATG13 and WIPI2 (Table 4). Importantly, cells deleted for ATG13 were also able to produce an LC3 response upon agonist stimulation (Suppl Fig 6A). These data argue that TRPML1 is not involved in canonical autophagy and are consistent with extensive recent data showing that the channel is likely involved in the non-canonical pathway termed CASM (conjugation of ATG8 to single membranes, (Goodwin et al., 2021; Lee et al., 2025). In support of this, overexpression of V5-tagged TRPML1 induced LC3 puncta formation even in normal growth conditions (Suppl Fig 6B) but did not affect WIPI2 localization (Suppl Fig 6C).

#### USP24

The deubiquitinase USP24 has been proposed to regulate autophagy via effects on ULK1 (Thayer et al., 2020). Additional siRNA experiments performed in stable HEK293 cells stably expressing GFP-LC3 but eliminated for ATG13 expression and stained for WIPI2 demonstrated that the LC3 and WIPI2 puncta phenotype seen in wild-type cells was not reproduced when canonical autophagy initiation was disabled (Suppl Fig 7). The absence of puncta induction in the ATG13-deficient context indicates that USP24 primarily regulates bulk autophagy rather than non-canonical pathways.

#### GLIPR2

GLIPR2 has been proposed to regulate autophagy via effects on the VPS34 complex (Zhao et al., 2021). Follow-up siRNA experiments in GFP-LC3 ATG13-knockout cells showed a partial reduction, but not elimination, of both LC3 and WIPI2 puncta compared with wild-type cells. This residual activity suggests that GLIPR2 influences both canonical and non-canonical pathways, consistent with a mixed mechanistic profile in autophagy regulation (data not shown).

#### ROCK2

ROCK2 has been proposed to regulate autophagy in cardiomyocytes (Shi et al., 2019). ROCK2 siRNA silencing in GFP-LC3–expressing cells and counterstained for phospho-TBK1 showed puncta formation containing both proteins but resembling aggregated material and not regular autophagosomes (Suppl Fig 8A). Importantly, ROCK2 silencing in GFP-ATG13 cells counterstained with WIPI2 did not reveal such structures indicating that this phenotype may be related to LC3 dynamics (Suppl Fig 8B). Finally, chemical inhibition of ROCK2 in GFP-LC3 expressing cells resulted in an increase in pTBK1-positive structures without accompanying GFP-LC3 puncta (Suppl Fig 8C). Given its possible function in LC3 dynamics, ROCK2 was further examined in iPSC-derived neuronal cells to derive a final conclusion on its potential function in autophagy (see below).

#### HUWE1

This E3 ligase was originally implicated in autophagy by the observation that it regulates expression of WIPI2, an important early autophagy protein that links formation of PI3P with lipidation of LC3 via its binding to ATG16 (Wan et al., 2018). Accordingly, downregulation of HUWE1 would prevent degradation of WIPI2 and should increase its levels. At the same time, a puzzling aspect of this hypothesis was our observation that simple overexpression of WIPI2 was not sufficient to activate autophagy in a variety of tissue culture cells (not shown). In our experiments, downregulation of HUWE1 with siRNA enhanced autophagy as seen by formation of WIPI2 puncta (Fig 5A-B), although it did not alter levels of WIPI2 either in basal conditions (Fig 5C) or following autophagy induction via inactivation of mTOR by PP242 (Fig 5D). On the other hand, one protein whose levels appeared to be altered upon HUWE1 downregulation was ULK1, the essential kinase of the early initiation machinery (Fig 5D, last blot). This increase was also seen for the phosphorylated form of ULK1 at serine 757 (Fig 5E, first 6 lanes). In cells deleted for ATG13, a binding partner of ULK1 in the ULK complex, the levels of ULK1 were reduced (as expected) but there was still an increase upon HUWE1 downregulation (Fig 5E last six lanes) suggesting that the effects of HUWE1 on ULK1 did not require the entire ULK complex. To examine the stability of ULK1 under normal conditions we used cycloheximide to block new protein synthesis and then assessed protein levels. We found that approximately 50% of the protein is degraded within 6 hours and the kinetics of this were similar in the presence or absence of HUWE1 (Fig 5F, G).

**Figure 5.**
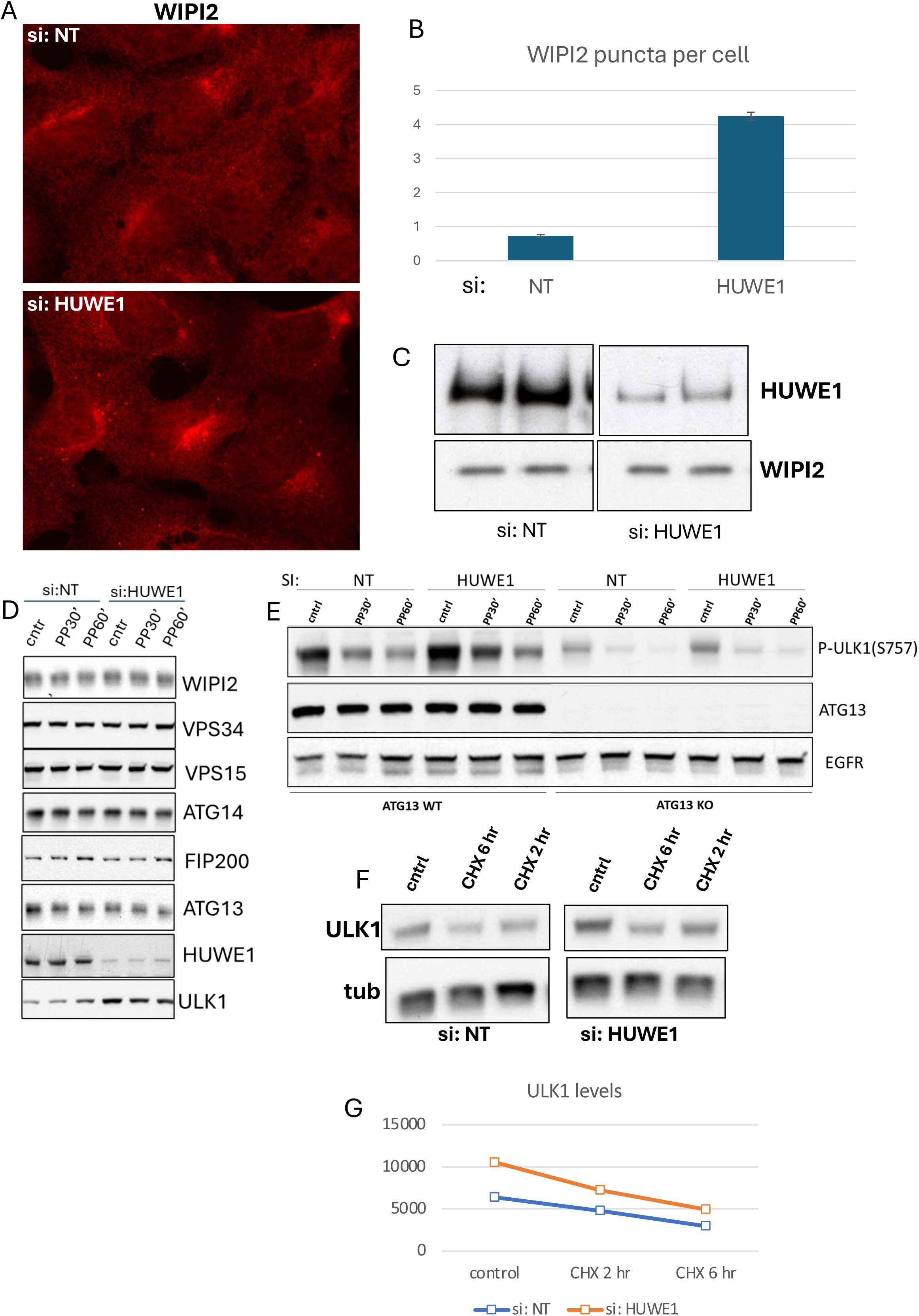
Characterization of HUWE1 as an autophagy regulator. A-C. HEK-293 cells were treated with siRNA against HUWE1 for 72 hrs and screened for formation of autophagy puncta using antibodies to endogenous WIPI2. Downregulation of HUWE1 routinely increased autophagic puncta in fed conditions by 3-4 fold. D. Cells treated with HUWE1 siRNA as above were incubated with mTOR inhibitor PP242 for 30’ or 60’ before lysis and immunoblotting for the indicated proteins. E. Parental HEK-293 cells or those deleted for ATG13 were treated with siRNA against HUWE1 and incubated with PP242 as shown in D. The lysates were immunoblotted for phospho-ULK1 (S757), ATG13 or EGFR as a loading control. F, G. HEK-293 cells treated with HUWE1 siRNA were incubated with cycloheximide for 2 or 6hr and then immunoblotted for ULK1. The rate of reduction of ULK1 is plotted in G.

Together, these mechanistic studies provide a refined view of the targets emerging from the screening cascade. ROCK2 displayed a TBK1 modulation with or without a concomitant effect on LC3 meriting further work. SYK, ARL8B and TRPML1 observations decrease the confidence in their modulation of the autophagic process. USP24 and GLIPR2 demonstrated reproducible autophagy phenotypes in HEK-293 cells and showed mechanistic features consistent with canonical or mixed-mode autophagy regulation. MAP2K5 produced autophagy-linked signals in screening assays but failed to validate robustly under orthogonal mechanistic conditions, supporting its inclusion as a lower-confidence follow-up target. HUWE1, in contrast, showed a consistent enhancement of early autophagy markers and mechanistically interpretable effects through ULK1 regulation, aligning with its strong performance in the screening cascade. On this basis, five targets (HUWE1, MAP2K5, USP24, GLIPR2, ROCK2) were selected for subsequent validation in neuronal autophagy assays, reflecting their mechanistic consistency, robustness across cell models, and relevance to distinct autophagy sub-pathways identified through the multi-screening, cross-platform strategy.

### Neuronal and in vivo Model Validation Identifies HUWE1 as an Autophagy Regulator for Further Target Development

To assess whether the regulatory effects observed in HeLa and HEK-293 cells translated into a neuronal context, we next evaluated the shortlisted autophagy-regulating targets - MAP2K5, ROCK2, HUWE1, USP24, and GLIPR2 - in human iPSC-derived neurons. Expression of these targets was downregulated using siRNA in iPSC-derived neuronal cells and, additionally, where available (for MAP2K5, ROCK2 and HUWE1), their function was inhibited by small molecules. Downregulation by siRNA was estimated by immunoblots as follows: MAP2K5, 93%; ROCK2, 62%; HUWE1, 70%; USP24, 81% and GLIPR2, 62%. The effects on autophagy, as measured by levels of LC3-II, for both siRNA and small molecules are shown in Fig 6. All five targets scored positive in the autophagy assays, confirming the reproducibility of our previous work in non-neuronal cell models. These effects validated the outcome of the multiplatform screening cascade and demonstrated that each candidate retained functional relevance in a cell type that is physiologically distinct and disease-relevant. Despite this broad validation, one target from this list (HUWE1) was chosen for immediate follow up in a *Drosophila* model, with the rest being judged to require further validation prior to progression. Our criterion for this decision was that the consistency and strength of autophagy induction varied among the targets across the range of experimental systems and the wish to select a target with the maximum potential to generate impactful new data. MAP2K5 and ROCK2 showed neuronal autophagy activation but had exhibited mechanistic ambiguity or partial inconsistency in earlier assays: for ROCK2 there were some concerns as to whether the autophagy enhancement is specific or due to wider effects on cellular physiology impacting autophagy consistent with no further increase of autophagy in siRNA-treated neurons upon bafilomycin treatment; for MAP2K5, as discussed above, an additional concern was whether its effects were connected to mTOR activation bringing the caveats of the multiple cellular changes when this kinase is inactivated. GLIPR2 was down-prioritised for immediate follow up due to a lack of chemical tools. For USP24, we noted a recent publication providing additional assessment of this target (Young et al., 2024) reducing the potential for our own studies to be impactful. Based on these considerations, these 4 targets were not prioritized owing to challenges identified by the consortium but remain of potential future interest.

**Figure 6.**
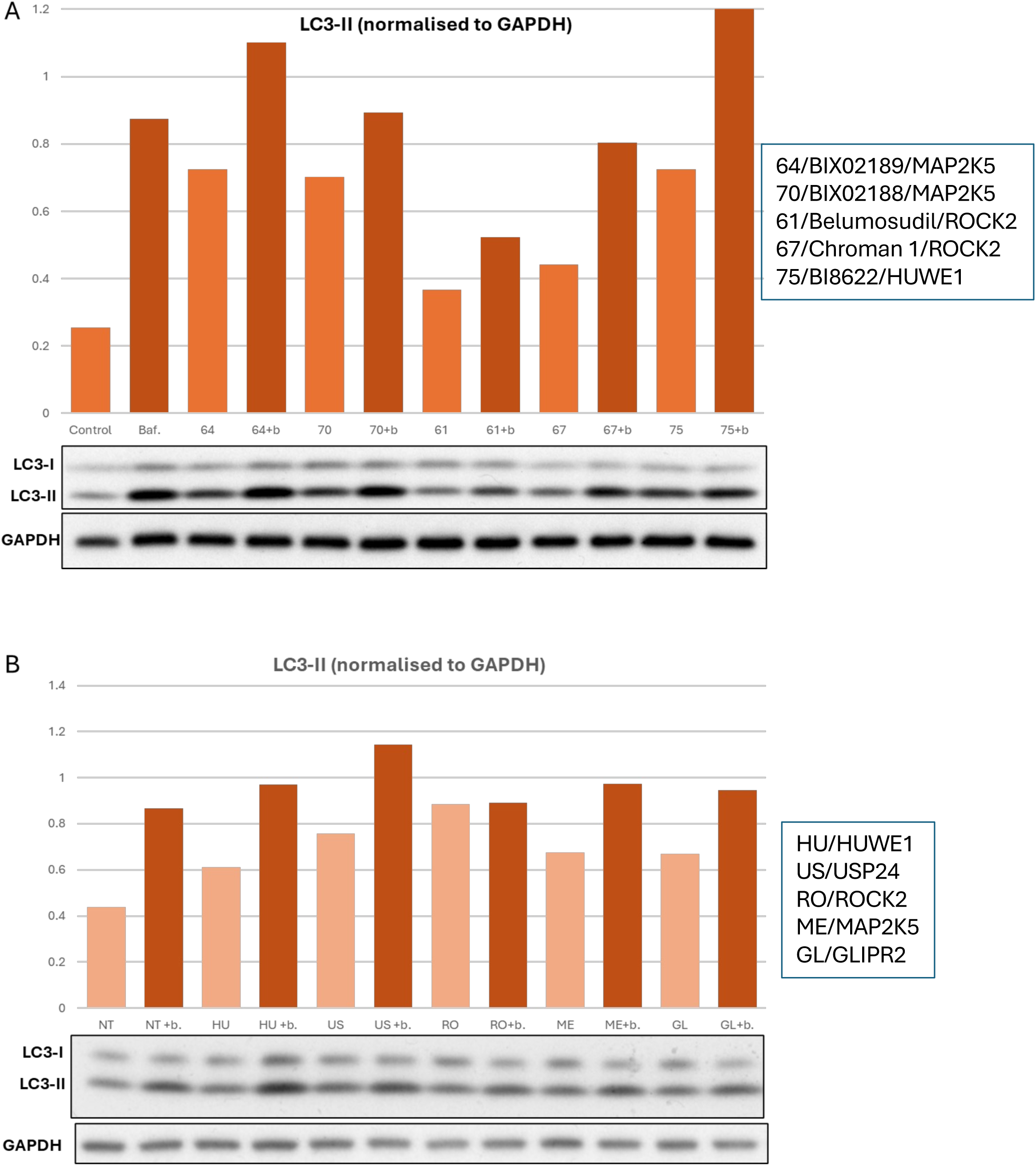
LC3-II formation in iPSC-derived neuronal cells upon pharmacological or genetic inhibition of various targets. A. Cells were treated with indicated compounds for 2 hr with or without bafilomycin A1 (indicated as Baf or b) and levels of LC3-II in the lysates were determined by immunoblotting as shown. B. Cells were treated with indicated siRNAs for 7 days and levels of LC3-II were determined as above. In both cases, intensity of the LC3-II signal was normalised to GAPDH used as loading control. These experiments were repeated 2 times with similar results.

In contrast, HUWE1 demonstrated clear, reproducible autophagy enhancement across all experimental modalities such as genetic and pharmacological inhibition, multiple imaging markers, analysis in both non-neuronal and neuronal models, and mechanistic assays showing a coherent effect on ULK1 levels and activity. This cross-platform robustness combined with its additional positive influence on huntingtin aggregate clearance in proteinopathy models distinguished HUWE1 as the most reliable negative-regulator of autophagy identified through the screening cascade.

In summary, the convergence of high-throughput imaging, mechanistic dissection, small-molecule modulation, and neuronal validation positioned HUWE1 as the strongest candidate for functional follow-up. Accordingly, HUWE1 was nominated as the top-priority target to progress into in vivo validation studies in *Drosophila* to test its relevance in whole-organism autophagy regulation and neurodegenerative disease models.

We turned to the well-established *Drosophila* model which is excellent for investigating conserved mechanisms of autophagy and neurodegeneration (Lorincz et al., 2017; Santarelli et al., 2023). Transgenic RNAi-mediated knockdown of the fly HUWE1 orthologue by two independent shRNA constructs led to a significant increase in mCherry-tagged Atg8a (the fly orthologue of LC3) in larval CNS neurons (Fig. 7A, B), consistent with elevated autophagy. Moreover, taking advantage of an established model of Huntington’s disease expressing an RFP-tagged N-terminal fragment of Htt containing 138 glutamines (RFP-Htt138Q, (Weiss et al., 2012), knockdown of HUWE1 significantly reduced the number of aggregates compared to control knockdown in larval neurons (Fig. 7C, D), consistent with increased degradation.

**Figure 7.**
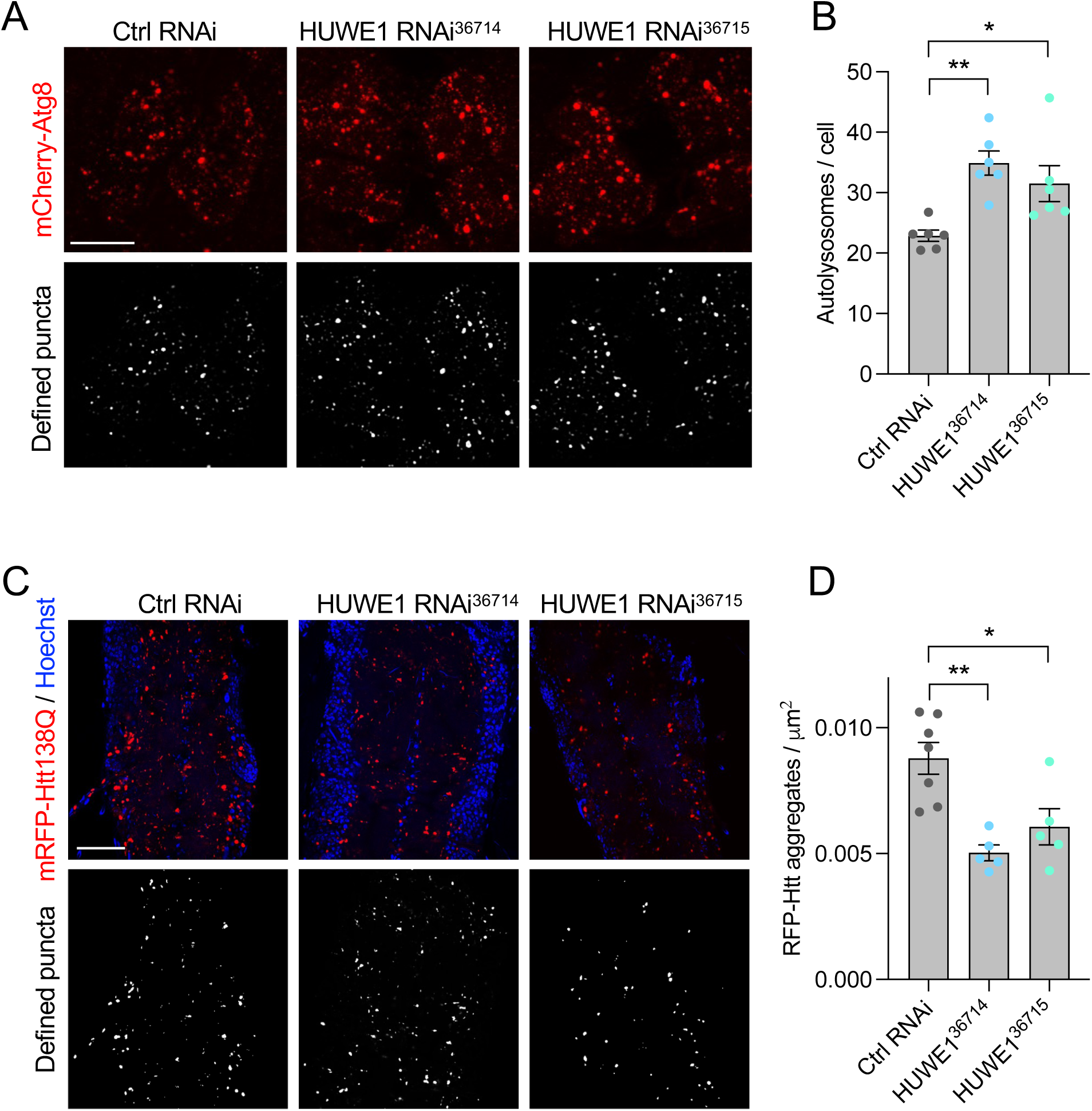
HUWE1 analysis in *Drosophila*. Genetic knockdown of HUWE1 in Drosophila shows increase in the levels of mCherry-ATG8 puncta (A, B) and a reduction in mRFP-Htt138Q aggregates in larval neurons (C, D). Charts show mean±SEM, n = 5-7 animals. Statistical analysis calculated by one-way ANOVA; * p<0.05, ** p<0.01.

In conclusion, each phase of our workflow informed the prioritization of targets for subsequent testing and guided the final selection as outlined in Table 5. Across the entire screening cascade from high-throughput imaging, through mechanistic profiling and small-molecule modulation, to validation in human neurons and whole-organism models, HUWE1 consistently emerged as the most convincing negative-regulator of autophagy with effects on huntingtin aggregate clearance, making it the candidate to advance with highest confidence into drug-discovery target validation for neurodegenerative diseases.

**Table 5.**
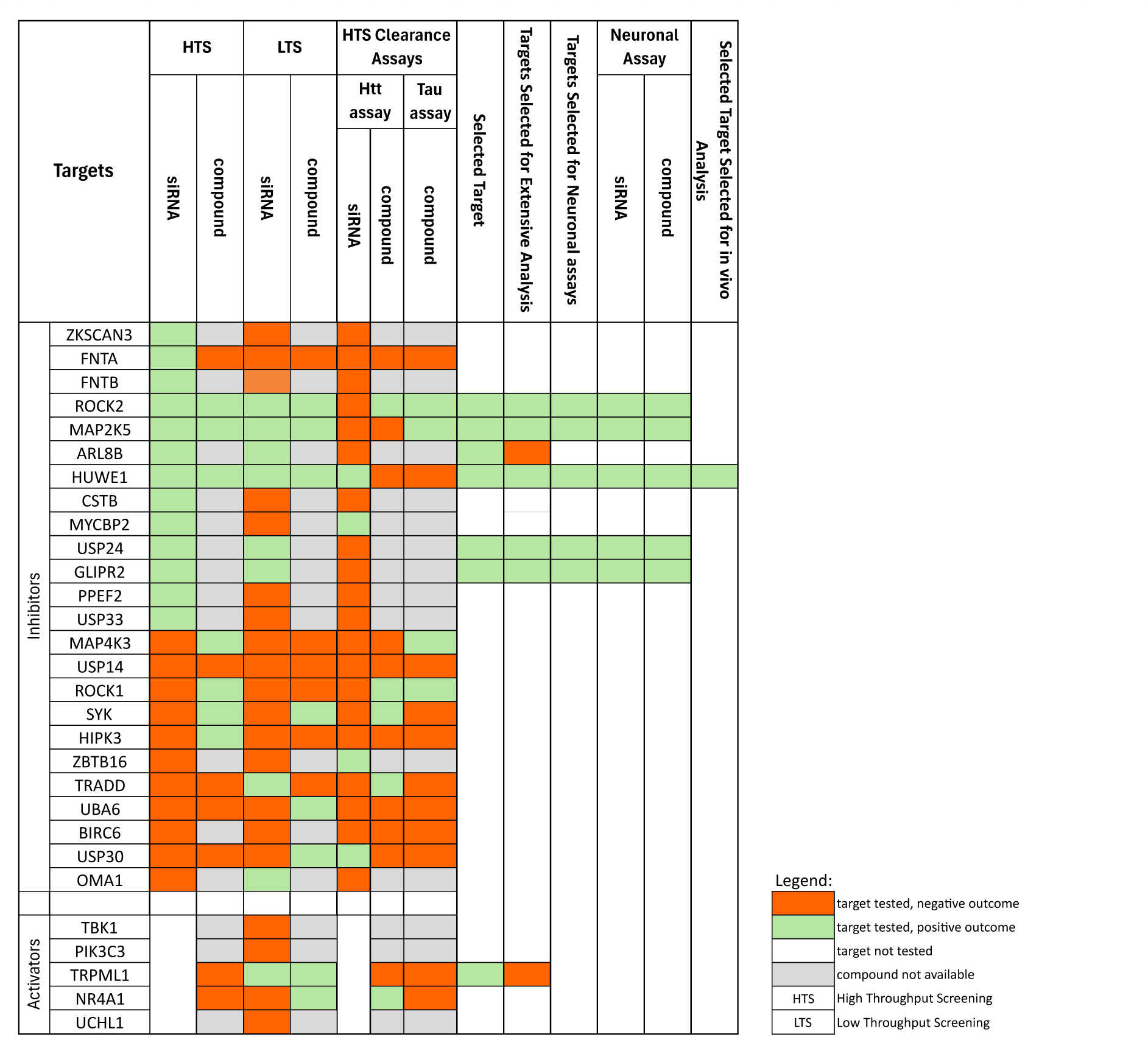
Details of the analysis for each autophagy target in the various assays are shown as well as their progression through the various experimental steps.

### Cross-platform Screening Reveals USP20 and USP33 as Targets Involved in Mitophagy

To systematically evaluate regulators of mitophagy, we implemented a high-throughput, image-based screen in ARPE-19 cells and assessed the impact of both genetic and pharmacological perturbations. ARPE-19 is a non-transformed human retinal epithelial line that stably expresses the mitophagy reporter tandem mCherry-GFP-MITO fusion protein targeted to the mitochondrial outer membrane via the FIS1(101-152) sequence (called mito-QC) (McWilliams et al., 2016). This reporter leverages the pH sensitivity of GFP, which is quenched upon lysosomal fusion, whereas mCherry remains stable, allowing specific identification of mitolysosomes as red-only puncta. From the initial candidate list, the consortium selected four genes - USP33, USP30, OMA1, and ROCK2 - based on prior reports of direct involvement in mitophagy regulation.

ARPE-19 cells responded robustly to the mitophagy inducer deferiprone (DFP), displaying a concentration- dependent increase in mitolysosome number that validated assay sensitivity and dynamic range (Fig. 8A, B). Gene silencing was achieved using SMARTpool siRNAs and yielded a knockdown efficiency of target suppression verified by qPCR that ranged from 75% up to over 90% (data not shown). Among the tested genes, ROCK2 and USP33 knockdown produced the strongest induction of mitophagy, increasing mitolysosome counts by approximately 5-fold and 7-fold, respectively, relative to non-targeting controls (Fig. 8C). Importantly, toxicity was monitored in all silencing conditions through nuclear counts, and no significant cell loss was observed compared to non-targeting controls. Pharmacological inhibition was also used to further explore the potential of these targets. Using an 8-point dose-response format (6.32 nM to 20 μM) and a 24-hour exposure window, we tested available inhibitors targeting ROCK2 (compounds Belumosudil and Chroman 1) and USP30 (a well-established regulator of mitophagy). Mitophagy scores were normalized to DMSO-treated cells as baseline (0) and DFP-treated cells as the maximal response (100). Toxicity was also assessed in parallel. The USP30 inhibitor induced only marginal increases in mitophagy (Fig. 8D). The lack of activity of the USP30 inhibitor is consistent with reports that inhibition of this target requires prior challenge with a mitochondrial toxin to trigger mitophagy in cellular assays (Fang et al., 2023). Whilst compound Chroman 1 (ROCK2 inhibitor) was inactive (Fig 8E), notably, compound Belumosudil elicited a modest but reproducible ∼20% increase in mitophagy activity, consistent with a possible role for ROCK2 in this pathway (Fig 8F). Toxicity was found to be minimal in all compound-treated conditions, with no reduction in cell viability distinguishable from DMSO controls. Overall, the siRNA and tool compound results validate USP33 and ROCK2 as functionally relevant mitophagy regulators.

**Figure 8.**
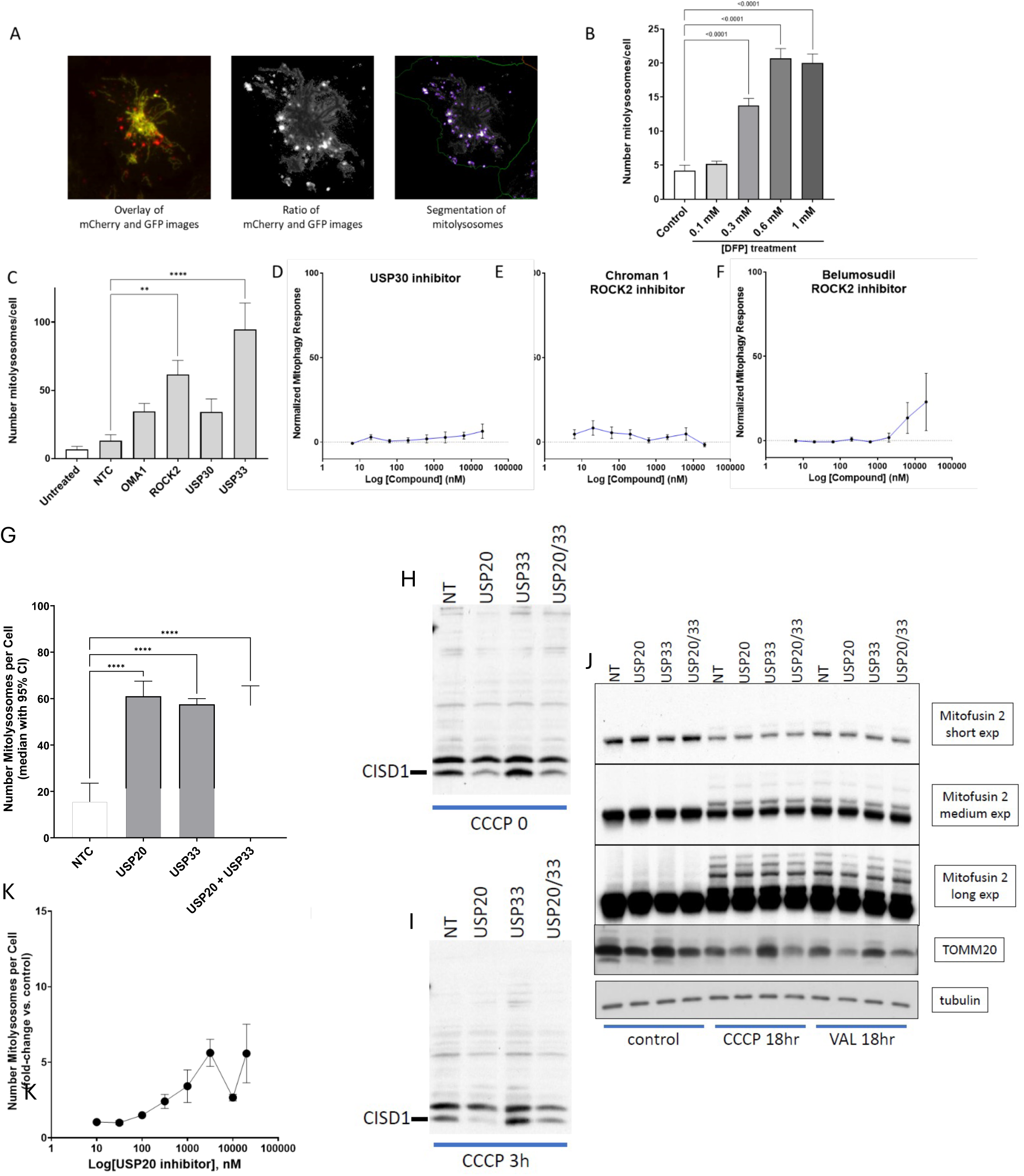
Characterization of USP20 and USP33 as mitophagy regulators. A. The mitophagy quantification pipeline. Left image: Overlay of mCherry and GFP channels showing the mitochondrial network (yellow) and mitolysosomes (red) after 24 h treatment with 1 mM deferiprone (DFP). Middle image: mCherry/GFP intensity ratio image highlighting regions with puncta corresponding to mitolysosomes. Right image: Segmentation mask of mitolysosomes created by thresholding pixels with mCherry intensity above the image average and mCherry/GFP ratio exceeding mitochondrial background. B. Dose-dependent treatment with DFP (0.1–1 mM) for 24 h, showing mitolysosome response. C. Mitophagy response in ARPE-19 cells with 12.5 nM siRNAs targeting indicated genes. Data represent three independent experiments with four technical replicates per condition, each consisting of 16 images acquired per well. Statistical significance was assessed by one-way ANOVA with Dunnett’s multiple comparisons test. ** p=0.0066; **** p<0.0001. Cell counts were monitored to evaluate toxicity. D-F. Concentration-response analysis of pharmacological inhibitors targeting candidate genes. Mitophagy activity was normalized with DMSO as 0 and DFP (0.8 mM) as 100. G. Mitophagy response upon siRNA downregulation of USP20, USP33 or USP20/33 in HEK-293 cells expressing the mito-QC reporter. Statistical significance was assessed by one-way ANOVA with Dunnett’s multiple comparisons test. **** p<0.0001. H, I. Reduction of mitochondrial protein CISD1 in HEK-293 cells downregulated for USP20 and USP33 as above in normal conditions or after treatment with CCCP for 3 hr as indicated. J. HEK-293 cells downregulated for USP20 and USP33 as above were treated with CCCP or Valinomycin for 18 hr and immunoblotted for the mitochondrial proteins Mitofusin 2 and TOMM20 as indicated. K. Dose response of mitophagy response upon treatment of ARPE-19 cells expressing mito-QC reporter with USP20 inhibitor at the indicated doses for 24 hr.

In parallel experiments mitophagy assays were carried using the same mitophagy reporter but expressed in HEK-293 cells. Given the positive result with USP33, we also evaluated knockdown of USP20, a protein that is highly homologous to USP33 and frequently works in tandem with it (Culver and Mariappan, 2021). Both USP20 and USP33 downregulation produced strong mitophagy responses in the HEK-293 cells (Fig 8G). The effects of USP20 and USP33 on mitophagy were also examined in biochemical assays measuring degradation of CISD1, TOMM20 and MFN2, all being implicated in this pathway (Martinez et al., 2024; Yoshii et al., 2011). In these assays, downregulation of USP20 produced a strong reduction in the levels of CISD1 and TOMM20, without strong effects on MFN2 (Fig 8H-J). The effect of downregulating both USP20 and USP33 together was similar to USP20 downregulation in those experiments. The mitophagy driven reduction in these proteins was seen under basal conditions with cells growing in normal medium as well as upon mitophagy induction with CCCP (Fig 8H, I). Finally, both siRNA-mediated depletion of USP20 and its pharmacological inhibition by GSK2643943A (Harrigan et al., 2018) showed mitophagy induction in ARPE-19 cells (Fig 8K).

In view of its possible role in mitophagy the localization of USP20 was investigated. Using antibodies whose signal was eliminated upon siRNA treatment against USP20 (not shown) we determined that the protein partially colocalizes with mitochondrial proteins such as TOMM20 and CIAP1 (Suppl Fig 9A) a pattern which was not altered upon mitophagy induction with CCCP. In other experiments we determined that USP20 also colocalises with markers of the endoplasmic reticulum (ER) (data not shown) suggesting that it may occupy the contact sites between mitochondria and ER. Finally, and consistent with previous reports, the depletion of USP20 increased the expression level of USP33 and vice versa (Suppl Fig 9B), suggesting that the total expression level of the two proteins may be controlled through a homeostatic mechanism.

In addition to USP20, USP33 and ROCK2, which scored positive in our mitophagy assays, we also looked at USP30, a well-established regulator of mitophagy. As described above, in our hands, chemical inhibition of USP30 did not induce mitophagy as measured by mitoQC, and siRNA against USP30 did not induce mitophagic structures. This is consistent with previous reports (Fang et al., 2023). Interestingly, in cells treated with the USP30 inhibitor we observed formation of ubiquitinated puncta which were positive for GFP-LC3 in cell lines stably expressing this reporter (Suppl Fig 10A). However, this was not seen in other cells lines stably expressing another autophagy reporter GFP-ATG13 (Suppl Fig 10B) or in parental cells (not shown). These ubiquitinated structures which were positive for LC3 in the GFP-LC3 expressing cells did not colocalise with mitochondria (Suppl Fig 10C) suggesting that they are not involved in mitophagy. Their provenance, and the reason why they appear upon GFP-LC3 (over)expression are unknown.

To translate the effects of USP20 or USP33 on mitophagy *in vivo*, we again turned to *Drosophila*. Transgenic lines expressing the mito-QC mitophagy reporter have been well-characterised (Lee et al, 2018). *Drosophila* encodes a single gene with equally high homology to both USP20 and USP33 and is hence named Usp20-33. Transgenic RNAi-mediated knockdown of Usp20-33 induced a significant increase in mitolysosomes in larval neurons, indicating upregulated mitophagy (Fig. 9). To address whether this increase occurred via parkin-mediated mitophagy we performed the same experiment in a *parkin* null mutant background. Loss of *parkin* did not diminish the Usp20-33 RNAi induced mitophagy (Fig. 9), demonstrating that this occurs via a parkin-independent mechanism.

**Figure 9.**
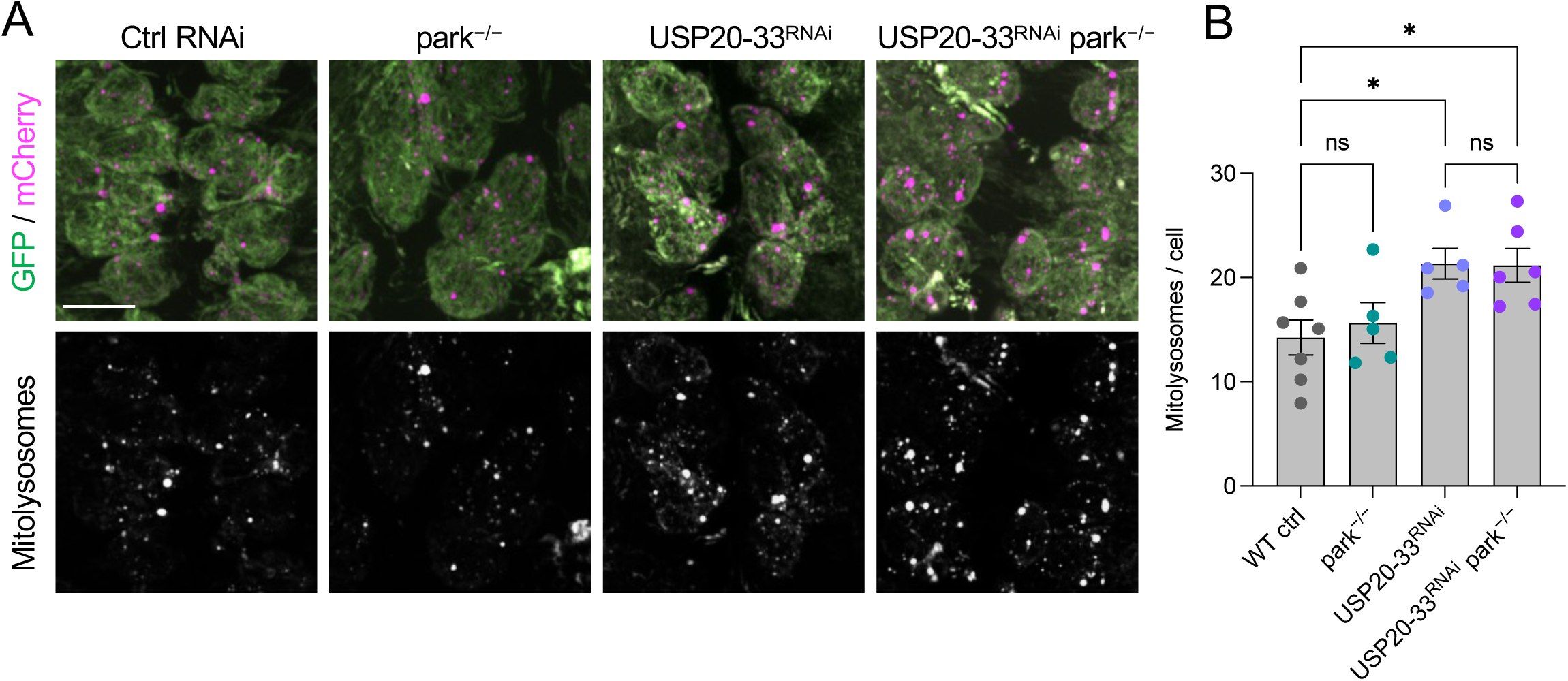
Usp20-33 analysis in *Drosophila*. Downregulation of Usp20-33 in Drosophila induces mitophagy (measured as appearance of mitolysosomes upon expression of a mito-QC reporter) which is not affected by loss of parkin. Chart in B shows mean±SEM, n = 5-8 animals. Statistical analysis calculated by one-way ANOVA; * p<0.05, ** p<0.01.

Together, these cross-platform cellular and in vivo data converge to identify USP20 and USP33 as the two most robust and reproducible modulators of mitophagy within our screening cascade. Given that mitochondrial dysfunction is a central pathogenic feature of multiple neurodegenerative diseases such as Parkinson’s disease, Alzheimer’s disease and Huntington’s disease, and that the enhancement of mitophagy represents a promising therapeutic strategy to counteract these deficits, USP20 and USP33 emerge as high confidence targets justifying further investigation within a drug-discovery framework specifically aimed at neurodegeneration.

## Discussion

Several whole-genome siRNA screens have been conducted in recent years to identify novel regulators of autophagy (Chan et al., 2007; DeJesus et al., 2016; Hale et al., 2016; Jung et al., 2017; Lipinski et al., 2010; McKnight et al., 2012; Orvedahl et al., 2011; Strohecker et al., 2015), and a number of screens using libraries of small molecules that affect autophagy have also been reported (Kuo et al., 2015; Mizushima and Murphy, 2020; Sarkar et al., 2007; Tian et al., 2011; Whitmarsh-Everiss and Laraia, 2021; Zhang et al., 2007). These efforts have provided some interesting and novel genes with moderate effects on autophagy and led to some useful tool compounds. Here we undertook a systematic analysis of reported autophagy regulators, using complementary assays and models to provide rigorous evidence supporting or negating the role of specific targets in autophagy. We included requirements that (a) their effects should be seen in multiple cell systems (including in iPSC-derived neurons) and in more than one laboratory and (b) subsequent examination of their function in a whole organism, namely *Drosophila*, should support initial findings in cell models. By combining these approaches and setting aims that go from single cells to whole organisms we set out to identify molecules of biological and therapeutic significance and also to build an understanding of the concordance between different orthogonal readouts of autophagy and protein clearance.

In the process of conducting our various small-scale screens we encountered the usual problems of hits that gave different results in the complementary assay systems run at our different laboratories. We found both biologically significant hits as well as those that produced statistically significant but biologically insignificant effects due to their very modest magnitude of change in the autophagic responses. Through the combined academic and industry expertise of the group, taking into account a number of considerations (reproducibility between labs, magnitude of effects, plausible mechanism, potential for pharmaceutical intervention down the line), the consortium selected three hits, HUWE1, and USP20/USP33, for analysis *in vivo*. The robustness of our validation strategy and triage and the importance of these hits was further supported by the observation that down-regulation of both hits in *Drosophila* produced results expected from our previous work: animals with reduced HUWE1 had increased autophagy and a modest but significant increase in aggregate clearance, whereas downregulation of USP20-33 increased mitophagy.

The mechanism of action of HUWE1 remains unclear and will become the subject of future work. We did not see an increase in WIPI2 protein levels upon HUWE1 siRNA-mediated downregulation as previously reported (Wan et al., 2018) whereas we saw a small but reproducible increase in the levels of ULK1, the only known protein kinase among the essential autophagy machinery (Mizushima, 2010). Interestingly, effects of HUWE1 depletion on ATG101, a component of the ULK complex together with ULK1, FIP200 and ATG13, were also reported (Lee et al., 2021) which would be consisted with our results focusing on ULK1. As part of our autophagy work over the last 15 years, we have generated many cell lines stably expressing autophagy proteins such as ATG13, VPS34, BECLIN1, LC3, WIPI2, ATG5, ATG7, ATG16, at levels higher than their endogenous counterparts and we have not seen with any of them a significant increase in basal autophagy. This indicates that this pathway is not likely to be regulated via increases in protein levels of essential autophagy proteins at least for mitotic cells in culture. The only exception to these findings is ULK1 whose expression has some unusual characteristics. On the one hand, it is not possible to derive a cell line stably overexpressing ULK1 presumably because its overexpression above a threshold is not tolerated, on the other hand, even modest transient overexpression of this protein is enough to induce increased autophagic activity (our unpublished data). This is consistent with work in *Drosophila* where it has been shown that animals overexpressing ATG1 (the homologue of mammalian ULK1) exhibit elevated autophagy in the absence of other signals (Scott et al., 2007). It is therefore plausible that HUWE1 could enhance autophagy by increasing levels of ULK1 and further work is required to clarify this point.

The pathway of mitophagy has been a major focus of interest amongst pharmaceutical companies, with the de-ubiquitinase USP30 being the most advanced and popular target being pursued (Antico et al., 2025; Bingol et al., 2014). Our work has provided some evidence supporting alternative targets among de-ubiquitinases - USP20 and its functional homologue USP33. Consistent with a role in mitophagy we found USP20 to be located partially on mitochondria and partially on the ER, with the latter being consistent with the reported roles of the protein in ERphagy (Zhang et al., 2024) and in the insertion of tail-anchored proteins on the ER (Culver and Mariappan, 2021). Interestingly, although the effects of USP20 depletion in our experiments were significant under basal conditions leading to reduction of the levels of a subset of mitochondrial proteins such as CISD1 and TOMM20, other mitochondrial proteins such as MFN2 were unaffected. This suggests that the effects of USP20 on mitophagy are specific for some proteins and future work should focus on explaining this observation. An additional important question related to USP20 is whether its function requires the PINK1/Parkin pathway or it engages with other ubiquitin-dependent machinery. Of note, others have also reported a role for USP33 in the regulation of PARKIN (Niu et al., 2020); our preliminary results in *Drosophila* suggests that its effects may be PINK1/Parkin-independent but more work is required to clarify this point. In addition, a recent study suggested that USP20 may have a function in stabilising PINK1 and in this way promote mitophagy induced by mitochondrial uncouplers such as CCCP (Park et al., 2025). Finally, an additional role for USP20 in the regulation of bulk autophagy has been reported due to effects on the stabilization of ULK1 (Kim et al., 2018). USP33 did not score positively in our bulk autophagy assays but this point is also worth exploring further.

In this study we have brought together the combined expertise of three academic and three industry teams to provide a rigorous re-assessment of the roles that 29 targets have in autophagy modulation. By optimizing analysis of target function using complementary assays and in vivo models, and interrogating results with both academic and drug discovery know-how, we identified 3 high-confidence targets for further validation. Importantly this resource has also highlighted the importance of such cross-platform approaches, in showing that several targets of previous focus in the literature do not show a robust role in autophagy regulation when evaluated across a wide range of functional readouts and biological systems. It is clear that there is no single definitive measure of autophagy induction, and it is only by following careful phenotypic and mechanistic characterisation that robust conclusions can be drawn. This resource therefore provides a comprehensive reassessment of the current literature around some autophagy targets, highlighting three high confidence targets for further study and provides an experimental toolkit for future validation studies.

On completion of the study the team reflected on the experience of working together. Throughout the project the consortium operated collaboratively, through regular steering meetings in which all members reviewed and discussed data and ideas and agreed objectives. As a team we found that this improved our decision-making and also served as an excellent opportunity for the different groups to learn from each other and understand each other’s approaches. The academic groups benefited from seeing directly the decision-making processes in industry that lead to target selection and the industrial groups benefited from seeing the data emerging from the academic laboratories and being able to contribute to experimental design and prioritisation. In conclusion all parties found this to be a positive experience and a constructive way to do science in a complex mechanistic area.

## Materials and Methods

### High-Throughput Autophagy Assay

#### siRNA Screening

HeLa cells stably expressing the tandem fluorescent autophagy marker mRFP-GFP-LC3 (Yoshimori T., Osaka University) were cultured in Dulbecco’s Modified Eagle Medium (GIBCO) supplemented with 10% fetal calf serum, 2 mM L-glutamine, 100 U/mL penicillin, 100 μg/mL streptomycin, and 0.6 mg/mL geneticin (G418). A high-throughput siRNA screen was performed in 384-well plates using the reverse transfection method with DharmaFECT 4 Transfection Reagent (Horizon Discovery) and ON-TARGETplus SMARTpool siRNAs (Horizon Discovery) at a final concentration of 25 nM. Control conditions included non-transfected cells, cells treated with GFP-targeting siRNA to assess transfection efficacy, and DharmaFECT 4-only controls.

After incubation with the siRNA-reagent complexes, 500 cells were seeded per well in a 384-well plate and cultured for 96 hours before fixation with formaldehyde and nuclear staining using 10 μg/mL Hoechst 33342 (ThermoFisher). The screening was carried out across two independent experiments, comprising a total of six plates with each gene tested in quadruplicate. Fluorescent images were acquired using the IN Cell Analyzer 6000 automated microscope, and image quantification was performed using CellProfiler. Cell-level data from each imaging channel were exported to Excel and analyzed using HC StratoMineR, with normalization of each plate to the median of non-targeting control wells.

#### Compound Screening

Autophagy-modulating compounds were screened using the same high-content imaging assay described for the siRNA screen. Compounds were tested in an eight-point dose-response format, ranging from 20 μM to 6.32 nM, and dispensed into 96-well plates using an Echo acoustic liquid handler. HeLa cells stably expressing mRFP-GFP-LC3 were seeded directly onto the compounds at a density of 5,000 cells per well for 48-hour incubations or 10,000 cells per well for 24-hour incubations. Following treatment, cells were fixed with 4% formaldehyde and stained with 10 μg/mL Hoechst 33342 (ThermoFisher) for nuclear labelling. The screening was conducted in four independent experiments, comprising a total of twenty 96-well plates, with each compound tested in a single replicate per concentration per plate. DMSO and 0.5 μM Torin 1 (an mTOR inhibitor) were included as reference controls on each plate, with DMSO values set to a baseline autophagy score of 0 and Torin 1 defining maximal autophagy induction at a score of 100.

Images were acquired using the IN Cell Analyzer 6000 automated fluorescence microscope. Image analysis was performed with CellProfiler, and single-cell data for all fluorescence channels were exported into Excel files, one per assay plate. These datasets were analysed using the web-based platform HC StratoMineR. In contrast to the siRNA screen, data normalization was performed using the normalized percent control method, where each measurement was scaled relative to the DMSO and Torin 1 control values from the same plate. Compound-induced cytotoxicity was assessed by quantifying the total number of Hoechst-positive nuclei per well, with significant reductions used as an indicator of toxicity.

Vendor details, catalogue numbers, and CAS identifiers for all compounds used in the screening are provided in the Suppl. Table 1.

#### Autophagy Score Calculation

Autophagy activity was quantified in HeLa cells expressing mRFP-GFP-LC3 that enables distinction between autophagosomes and autolysosomes based on pH-sensitive GFP quenching. Autophagosomes were identified as puncta positive for both GFP and RFP, while autolysosomes—formed after fusion with lysosomes—appeared as RFP-only puncta due to GFP quenching in the acidic environment. Four image-derived features were extracted per cell to characterize the autophagic response: (i) average GFP intensity (reflecting LC3 expression), (ii) average number of GFP+ (autophagosome) puncta per cell, (iii) average number of RFP+ (autolysosome) puncta per cell, and (iv) the GFP/RFP fluorescence intensity ratio within RFP+ puncta, serving as an indicator of autophagic flux. Each feature was normalized individually and an integrated autophagy score was computed as the average of the four normalized features, capturing both the abundance and dynamic flux of the autophagy pathway.

### High-Throughput GFP-HttQ74 Clearance Assay

#### siRNA Screening

A high-throughput imaging-based clearance assay was developed to assess the degradation of mutant huntingtin in PC12 cells stably expressing a doxycycline-inducible GFP-tagged exon 1 of huntingtin containing a 74-glutamine repeat (GFP-HttQ74, a kind gift from Prof David Rubinsztein). Cells were maintained in RPMI 1640 medium supplemented with 10% horse serum, 5% fetal bovine serum, 2 mM L-glutamine, 100 U/mL penicillin, 100 μg/mL streptomycin, 0.1 mg/mL geneticin (G418), and 0.07 mg/mL hygromycin. For siRNA screening, 3,000 cells were seeded per well in collagen-coated 384-well plates using reverse transfection with DharmaFECT 4 (Horizon Discovery) and 12.5 nM ON-TARGETplus SMARTpool siRNAs (Horizon Discovery). For HUWE1, a pool of four individual predesigned rat siRNAs (Merck) was used. After 19 hours, doxycycline was added to induce GFP-HttQ74 expression. Following a 96-hour incubation, cells were fixed with 4% formaldehyde and stained with 10 μg/mL Hoechst 33342 (ThermoFisher) to label nuclei. Fluorescent images were acquired using the IN Cell Analyzer 6000 automated microscope, and image analysis was performed with CellProfiler. Cell-level data from all channels were exported to Excel files (one per plate), integrated in KNIME, and subsequently uploaded to the web-based platform HC StratoMineR for high-content analysis. Data normalization was performed using the normalized percent control method, where uninduced cells (no huntingtin expression) represented 100% clearance and induced cells treated with non-targeting control siRNA defined 0% clearance. Normalization was carried out on a per-plate basis.

#### Compound Screening

Autophagy-dependent clearance of GFP-HttQ74 was assessed in a compound screen using the same imaging platform as described above. Compounds were solubilized in DMSO and dispensed in an eight-point dose-response format (20 μM to 6.32 nM) into collagen-coated 96-well imaging plates. PC12 cells were seeded at a density suitable for a 48-hour incubation directly onto compound-containing wells in the presence of doxycycline to induce GFP-HttQ74 expression (uninduced controls were included on each plate). Following treatment, cells were fixed with 4% formaldehyde and stained with 10 μg/mL Hoechst 33342. As with the siRNA screen, images were acquired using the IN Cell Analyzer 6000 and analyzed using CellProfiler. Single-cell fluorescence data were exported to Excel, processed in KNIME, and analyzed using HC StratoMineR. Clearance data were normalized using the normalized percent control method, where uninduced cells defined 100% clearance and induced DMSO-treated cells defined 0% clearance.

Compound-associated toxicity was calculated as the reduction in the number of Hoechst-positive nuclei, with 100% toxicity representing complete loss of cells and 0% toxicity indicating no change from DMSO controls. A four-parameter logistic regression model was used to determine EC₅₀ values for both clearance and toxicity profiles.

#### Clearance Score Calculation

Clearance activity was quantified using a composite metric termed the “clearance score,” which reflects both the reduction of soluble huntingtin and the elimination of aggregates. Two image-derived features were extracted per cell: (i) average cellular GFP intensity, representing the soluble GFP-HttQ74 pool, and (ii) the percentage of cells containing at least one aggregate, based on detection of bright GFP-positive puncta. Each feature was normalized individually, and the clearance score was computed as the average of the two normalized values, providing an integrated readout of the efficacy of huntingtin clearance.

### High-Throughput HiBiT-Based Tau Clearance Assay

A luminescence-based high-throughput assay was developed to assess compound-induced clearance of tau protein in HeLa cells transiently transfected with a plasmid encoding the human 0N4R Tau isoform carrying the P301S mutation and fused to an 11-amino acid HiBiT tag (0N4R-Tau-HiBiT^P301S). The HiBiT tag enables sensitive quantification of intracellular tau levels via the Nano-Glo® HiBiT Lytic Detection System (Promega #N3040), which provides luminescent signal upon complementation of HiBiT with the exogenously added LgBiT polypeptide. HeLa cells were seeded and transfected using Lipofectamine 3000 (ThermoFisher) according to the manufacturer’s instructions. After a 6-hour transfection period, cells were trypsinized and transferred to 384-well plates at 3,000 cells per well containing pre-dispensed compounds in a nine-point dose-response format ranging from 53.1 µM to 6.32 nM. Following a 24-hour incubation, Nano-Glo® HiBiT Lytic Detection reagents were added directly to each well, and luminescence was measured using a plate reader. Tau clearance activity was quantified by normalizing luminescence values using the normalized percent control method, where untransfected cells defined 100% clearance and DMSO-treated transfected cells defined 0% clearance on a per-plate basis. In parallel, compound-associated cytotoxicity was assessed using the CellTiter-Glo® Luminescent Cell Viability Assay (Promega #G7571), which quantifies ATP as an indicator of metabolically active cells. Luminescence signals from both assays were fitted using a four-parameter logistic regression model to calculate EC₅₀ values for tau clearance and toxicity, respectively.

### Mitophagy ‘Mito-QC’ Assay

Mitophagy was assessed using a high-content imaging assay in ARPE-19 cells (a human retinal pigment epithelial cell line) stably expressing a dual-fluorescent mitophagy reporter construct, in which mCherry and GFP are fused to the mitochondrial outer membrane localization sequence (amino acids 101–152) of FIS1 (a kind gift from Prof Ian Ganley) (Allen et al., 2013; McWilliams et al., 2016). This reporter exploits the pH sensitivity of GFP, which is quenched in the acidic lysosomal environment, whereas mCherry remains fluorescent, allowing distinction between mitochondria localized in the cytoplasm (GFP⁺/mCherry⁺) and those undergoing mitophagy (mCherry⁺ only). Mitophagy was also assessed in HEK-293 cells stably expressing the Mito-QC reporter as above.

#### siRNA screening

ARPE-19 cells were reverse transfected in 384-well plates at 175 cells per well using 12.5 nM Dharmacon™ SMARTpool siRNAs and DharmaFECT 4 transfection reagent. Non-targeting and GFP-targeting siRNAs were included as controls. After a 96-hour incubation, cells were fixed using 3.7% formaldehyde in PBS supplemented with 200 mM HEPES (pH 7.0) and 10 μg/mL Hoechst 33342 for 15 minutes at room temperature. Following washing, cells were further incubated in PBS containing 10 μg/mL Hoechst 33342 and Alexa Fluor™ 647-conjugated Wheat Germ Agglutinin (Invitrogen #W32466) for 80 minutes at room temperature.

#### Compound screening

DMSO-solubilized compounds were dispensed into 384-well plates in a nine-point dose-response format ranging from 20 μM to 6.32 nM. ARPE-19 cells were seeded directly onto compound-containing wells and incubated for 24 hours. Fixation and staining were performed as described for the siRNA screen.

DMSO and 1 mM Deferiprone (DFP), a known mitophagy inducer, were included as controls in both siRNA and compound screening.

#### Image Analysis

Images were acquired using the IN Cell Analyzer 6000 automated fluorescence microscope. Image analysis was carried out in CellProfiler, and single-cell data for each fluorescence channel were exported to Excel and processed using KNIME before being uploaded to Dotmatics for downstream analysis. Data normalization was performed using the normalized percent control method, where DMSO-treated cells defined a mitophagy score of 0 and DFP-treated cells defined a score of 100. Cytotoxicity was assessed by quantifying Hoechst-positive nuclei, with 0% toxicity indicating no loss compared to DMSO controls and 100% toxicity indicating complete cell loss.

To detect mitophagy events, image processing was based on ratiometric analysis of mCherry and GFP signals. First, images were filtered using a median filter to reduce background noise. Cells were segmented based on Hoechst and WGA staining. A pixel-wise ratio image was generated by dividing the mCherry signal by the corresponding GFP signal. Thresholding was applied using two parameters: (i) the average mCherry/GFP intensity ratio in untreated cells, representing baseline mitochondrial localization, and (ii) the average mCherry intensity to exclude regions with low signal that may yield false-positive ratios. A binary mask was then applied to isolate high confidence mitolysosomes by exploiting the loss of GFP fluorescence in lysosomes, and mitophagy was quantified by identifying and counting these objects in the ratio image.

### Preparation of iPSC-derived neuronal cells

KOLFC1 iPS cells (a kind gift from Dr Bill Skarnes) were induced to generate neuronal stem cells which were subsequently differentiated into early neurons and then matured into mature neurons, of primarily cortical lineage. Typically, the iPSCs were grown for 6 days, induced and cultured for 17 days, differentiated for 5 days and matured from 7 to 30 days depending on the experiment. All media used were originally from Axol Bioscience but in subsequent experiments from StemCell Technologies following their exact protocols.

### Immunofluorescence

Microscopy Protocols were described recently (Karanasios and Ktistakis, 2015; Zachari et al., 2019). Briefly, cells on coverslips were fixed in 3.7% Formaldehyde, permeabilized in 0.1% NP40 and stained in a blocking solution containing fish gelatin and 0.05% NP40. Antibodies used and their final dilution are as follows: mouse anti-WIPI2 (Bio-Rad), 1:200; rabbit anti-phospho(S172)-TBK1 (Cell Signalling), 1:50; rabbit anti-FIP200 (ProteinTech), 1:100; mouse anti-TOMM20 (Abcam), 1:100; rabbit anti-ATG13 (Sigma), 1:100; rabbit anti-LC3 (Sigma) mouse anti-Ubiquitin FK2 (Enzo). Secondary antibodies were from Jackson ImmunoResearch of appropriate species provenance and coupled with Cy2 or Cy3 dyes.

### Lysates and Immunoblots

Cells were lysed on ice in appropriate volume of lysis buffer [50 mM Tris pH 8.0, 50 mM KCl, 1mMEDTA pH8. 0,1% IGEPAL, 0.6mM PMSF, Complete Mini, EDTA-free tablet (Roche)] supplemented with 10 % 50mM NaF, 0.1% 10mg/ml leupeptin and 0.5% 0.2M Sodium orthovanadate. Lysates were collected by scraping and centrifuged for 10 min at 14,000 rpm at 4C. The supernatant was collected in fresh tubes on ice. Protein concentration of the lysates was determined using the BCA protein assay kit (Thermo Scientific Pearce). For electrophoresis, samples were combined with 2 x Laemmli sample buffer [20mM Tris-Cl pH 6.8, 2% sodium dodecyl sulphate (SDS), 10% glycerol, 0.1M dithiothreitol (DTT)] in a 1:1 ratio. Samples were then heated for 90 seconds at 95 °C before loading. Gels were wet transferred overnight to 0.45mm Immobilon-P transfer membranes (Millipore). Incubations with primary and secondary antibodies, and signal development using ECL (ECL Western Blotting Detection Reagent, GE Healthcare/Amersham Biosciences) followed standard protocols. For immunoblots, the following antibodies at the indicated dilutions were used: rabbit anti-ATG9 (Cell Signalling), 1:1500; rabbit anti-ATG13 (Sigma), 1:1000; rabbit anti-FIP200 (ProteinTech), 1:1000; mouse anti-GAPDH (Biogenesis), 1:100,000; rabbit anti-phospho(S172) TBK1 (Cell Signalling), 1:1000; rabbit anti-ULK1 (ProteinTech), 1:1000; rabbit anti-phospho(S757) ULK1 (Cell Signalling) 1:1000, mouse anti-TBK1 (Santa Cruz), 1:500.

### siRNA Experiments

Cells were seeded in 6-well plates to reach 70%–80% confluency 24 h later. Transfections were performed using DharmaFect (Dharmacon) with SmartPool siRNA oligos against non-targetting and target specific oligos (Dharmacon) for 72 hr. In some experiments it was necessary to perform double downregulation as follows: Cells were plated in the morning and transfected for the first time in the afternoon with 80 pmol of each siRNA SmartPool. The cells were re-transfected two days later with the same amount of siRNA and examined for the various assays two days later.

### siRNA experiments in iPSC-derived neuronal cells

Neuronal cells at approximately 7 days after differentiation (a time point that allows freezing and thawing to maintain consistent starting pools) were allowed to recover for 3 days in BrainPhys neuronal medium (StemCell Technologies) and were then switched to Neurobasal medium for the siRNA protocol. Neuronal cells were incubated with Accell siRNA (Dharmacon) following exactly the manufacturer’s instructions.

Effective downregulation of the various gene targets was determined in preliminary experiments, and it ranged from 6 to 8 days of incubation with the siRNA.

### Quantitation and statistical analysis

For quantitating puncta formation by immunofluorescence, 10 images (technical repeats) were randomly acquired for each treatment condition and quantified using the ImageJ cell counter plugin. Data from biological repeats were combined, log-transformed, and a two-way ANOVA was performed to obtain statistical significance and finally plotted as Mean±SD. All statistical analysis was checked by Dr Anne Segonds-Pichon, the statistician of the Babraham Institute.

#### Drosophila stocks and husbandry

Flies were raised under standard conditions in a temperature-controlled 12:12 hour light/dark cycle incubator at 25 °C and 65% relative humidity, on standard *Drosophila* food containing cornmeal, agar, molasses, yeast and propionic acid (Department of Genetics, University of Cambridge). Transgene expression was induced using the pan-neuronal driver *nSyb-*GAL4 driver except for expressing UAS-Htt138Q for which the weaker pan-neuronal *elav-*GAL4 driver was used. The following strains were obtained from Bloomington *Drosophila* Stock Center (BDSC): *w*^1118^ (BDSC_6326), *elav-*GAL4 (BDSC_8765), *nSyb-*GAL4 (BDSC_51635), *UASp-mCherry-Atg8a* (BDSC_37750), *UAS-mito-QC* reporter (BDSC_91640), *UAS-luciferase RNAi* (BDSC_31603), *UAS-HUWE1-RNAi* (BDSC_36714, BDSC_36715), *UAS-Usp20-33-RNAi* (VDRC_v110250), *park^25^* (BDSC_95259) (Greene et al, 2003). The *UAS-HTT.138Q.mRFP* line was a kind gift from Prof. Troy Littleton.

#### Drosophila Microscopy

Fluorescence imaging was conducted using a Zeiss LSM 880 confocal microscope (Carl Zeiss MicroImaging) equipped with Nikon Plan-Apochromat 63x/1.4 NA oil immersion objectives. For mito-QC imaging, the Andor Dragonfly spinning disk microscope was used, equipped with a Nikon Plan-Apochromat 100x/1.45 NA oil immersion objective and iXon camera. Z-stacks were acquired with 0.2 μm steps.

### Quantification and analysis methods

A single image-stack was generated per animal in which an area of interest was selected by choosing 6-10 cells per image. The quantification of autolysosomes and mRFP-Htt138Q puncta was performed using FIJI (Image J) with the 3D Objects Counter Plugin. The threshold was based on matching the mask with the fluorescence. A minimum size threshold of 0.05 μm^3^ was set to select autolysosomes. The quantification of mitolysosomes was performed as described in (Lee et al, 2018) using Imaris (version 9.0.2) analysis software. Briefly, a rendered 3D surface was generated corresponding to the mitochondrial network (GFP only). This surface was subtracted from the mCherry signal which overlapped with the GFP-labelled mitochondrial network, defining the red-only mitolysosomes puncta with an estimated size of 0.5 μm and a minimum size cut-off of 0.2 μm diameter determined by Imaris.

## Resource Availability

Requests for further information and resources should be directed to and will be fulfilled by the lead contact, John Skidmore.

Plasmids:

- Plasmid encoding the human 0N4R Tau isoform carrying

the P301S mutation and fused to an 11-amino acid HiBiT tag (0N4R-Tau-HiBiT^P301S).

## Supplementary Figures

**Suppl Figure 1. Deconvolution of siRNA pools.** A-B. Validation of siRNA pool hits by deconvolution. Each gene for which the SMARTpool increased autophagy was further tested with its four individual siRNAs. Genes were retained as autophagy modulators if at least 3 of 4 siRNAs reproduced the phenotype. C. Assessment of cell viability based on nuclear staining (Hoechst). Relative cell count was determined for each siRNA condition.

All data are from two independent experiments, each performed across four plates, with quadruplicate wells per condition. Each plate included GFP Duplex I control siRNA to confirm transfection efficiency through reporter gene knockdown (nearly 100% expression inhibition, not shown). Statistical significance was determined using ordinary one-way ANOVA followed by Dunnett’s post hoc test, comparing each condition to the non-targeting control. p < 0.05 was considered significant (*). Ns = non-significant. Values are expressed as mean ± SEM.

**Suppl Figure 2. Clearance assays for GFP-Htt(Q74) and Tau(P301S)-HiBiT.** A. Genetic modulation of autophagy-related targets in the GFP-Htt aggregate clearance assay. PC12 cells stably expressing doxycycline-inducible GFP-Htt(Q74) were treated with siRNAs targeting candidate genes. Following transfection, aggregates were induced by doxycycline for 96 hours. “Clearance Score” combined mean GFP intensity (reflecting soluble Htt) and the proportion of cells with ≥1 visible aggregate. It was scaled between 0 (induced, non-targeting siRNA control) and 100 (uninduced, full clearance). Positive Control: mTOR knockdown; negative Control: ATG13 knockdown. Data are mean ± SEM from at least 2 independent experiments, each performed in 3 technical replicates. Statistical analysis was conducted using ordinary one-way ANOVA with Dunnett’s multiple comparisons test (* = p < 0.05 compared to non-targeting control). B. Compound-based modulation of target activity in the GFP-Htt aggregate clearance assay. Compounds targeting the candidate genes were tested in a 48-hour incubation in the presence of doxycycline. Clearance scores were calculated as in (A), and normalized between 0 (DMSO-treated, induced control) and 100 (uninduced control). For each compound, two dose-response curves were generated: one for aggregate clearance and one for cell viability. Data are representative of two independent experiments each performed with one plate with quadruplicate wells per condition. C. Compound-based modulation of tau clearance using a 0N4R-Tau-HiBiT(P301S) assay in HeLa cells. Cells were transfected with a plasmid encoding the aggregation-prone 0N4R-Tau(P301S) isoform tagged with a HiBiT peptide. Six hours post-transfection, cells were replated onto compound-loaded 96-well plates for 24 hrs. Tau levels were quantified by luminescence. Clearance scores were normalized between 0% clearance (DMSO-treated transfected control) and 100% clearance (untransfected control). As in (B), two dose-response curves were generated per compound. EC₅₀ values for both parameters were calculated and plotted to evaluate clearance efficacy relative to compound toxicity. Data are representative of two independent experiments; each performed with one plate and four technical replicates per condition.

**Suppl Figure 3. Characterization of SYK as an autophagy modulator.** A. HEK-293 Cells were treated with the SYK inhibitor GSK143 as indicated, in the presence or absence of Bafilomycin A1 and levels of LC3-II were determined by immunoblotting. B. Similar analysis as in A but LC3 puncta were examined by immunofluorescence. C. Cells were treated with SYK siRNA and the levels of LC3-II following 4-way analysis (control/PP242/Bafilomycin A1/PP242+Bafilomycin A1) to measure autophagic flux were determined by immunoblotting.

**Suppl Figure 4. Characterization of MAP2K5 as an autophagy modulator.** A. HEK-293 cells were treated with two MEK5 inhibitors (BIX02188 and BIX02189) for 1 hr and were then stained with antibodies to endogenous WIPI2 and FIP200. B. Similar experiment as in A but using several time points of treatment was quantitated for WIPI2 and FIP200 colocalizing puncta. C. Cells were treated with the two MEK5 inhibitors or with the mTOR inhibitor PP242 or were starved of amino acids for 1 hr as shown, lysed and the lysates immunoblotted for phospho-P70 and phospho-P85, for P70 and P85 or for phospho-ULK1 (S757) as shown.

**Suppl Figure 5. Characterization of ARL8 as an autophagy modulator.** HEK-293 cells were treated with siRNA against ARL8 for 72 hr and then a 4-way analysis for autophagic flux was performed (control/PP242/bafilomycin A1/PP242+bafilomycin A1) assaying levels of LC3-II by immunoblotting. The intensity of the LC3-II bands normalised to GAPDH are shown in the graph and correspond to the lanes above.

**Suppl Figure 6. Characterization of TRPML1 as an autophagy modulator.** A. Number of LC3 puncta per cell was determined in parental HEK-293 cells or in those deleted for ATG13 as indicated using 4 different TRPML1 agonists at 10 μM. The levels of LC3 puncta were significantly elevated following those treatments but were not reduced in cells devoid of ATG13, indicating that their occurrence maps to the non-canonical autophagy pathway termed CASM. B. Elevated LC3 puncta were also seen in cells over-expressing TRPML1 tagged with V5. C. LC3 puncta elevated by treatment with the agonist ML-SA1 did not colocalize with WIPI2, indicating that they are not related to autophagy induction.

**Suppl Figure 7. Characterization of USP24 as an autophagy modulator.** USP24 was downregulated by siRNA in HEK293 cells stably expressing GFP-LC3 or in HEK293 cells stably expression GFP-LC3 but eliminated for the expression of ATG13 as indicated. In both cases, cells were counterstained with antibodies against endogenous WIPI2.

**Suppl Figure 8. Characterization of ROCK2 as an autophagy modulator**. A. HEK293 cells stably expressing GFP-LC3 were treated with siRNA against ROCK2 and counterstained with antibodies against endogenous phospho-TBK1. B. HEK293 cells stably expressing GFP-ATG13 were treated with siRNA against ROCK2 and counterstained with antibodies against endogenous WIPI2. C. HEK293 cells stably expressing GFP-LC3 were treated with the ROCK2 inhibitor Belumosudil SLx-2119 for 18 hr and counterstained with antibodies against endogenous phospho-TBK1.

**Suppl Figure 9. Characterization of USP20 and USP33**. A. HEK-293 cells were left untreated or treated with CCCP for 4hr before fixation and indirect immunofluorescence using antibodies specific to USP20 and two mitochondrial proteins, TOMM20 and CIAP1 as shown. Independently of CCCP-induced mitochondrial fragmentation, a portion of USP20 co-localizes with mitochondria. B. A variety of cell lines as shown were treated with siRNA against USP20 or USP33 and the relative levels of the two proteins were determined by immunoblotting. Reduction of USP20 induces a large increase in the levels of USP33 whereas in the opposite scenario the effect on USP20 when USP33 is reduced is more modest.

**Suppl Figure 10. Cellular effects of USP30 inhibition.** A. HEK-293 cells stably expressing GFP-LC3 were treated with the USP30 inhibitor for 60 min as shown and stained for immunofluorescence with antibodies to ubiquitin. Puncta stained for ubiquitin and LC3 are induced by the drug. B. HEK-293 cells stably expressing GFP-ATG13 were treated with USP30 inhibitor as above and stained for ubiquitin. In those cells no ubiquitin puncta are induced. C. In HEK-293 cells stably expressing GFP-LC3 the LC3 puncta induced by USP30 inhibition do not colocalise with the mitochondrial protein TOMM20.

**Suppl Table 1.** Vendor information for compounds used in the screening. The table lists, for each compound, the supplier, catalogue (vendor) number and CAS registry number.

| Mode | Compound | Target | Vendor | Vendor ID | CAS |
| --- | --- | --- | --- | --- | --- |
| Inhibitors | Neratinib | MAP4K3 | MedChemExpress | HY-32721 | 698387-09-6 |
|  | Tipifarnib | FNTA | Tocris Bioscience | 6410 | 192185-72-1 |
|  | IU1 | USP14 | Cayman Chemical | CAY10617 | 314245-33-5 |
|  | Azaindole 1 | ROCK1 | MedChemExpress | HY-10319 | 867017-68-3 |
|  | GSK269962A |  | MedChemExpress | HY-15556 | 850664-21-0 |
|  | Chroman 1 | ROCK2 | Tocris Bioscience | 7163 | 1273579-40-0 |
|  | Belumosudil |  | MedChemExpress | HY-15307 | 911417-87-3 |
|  | Entospletinib | SYK | MedChemExpress | HY-15968 | 1229208-44-9 |
|  | GSK143 |  | Tocris Bioscience | 6362 | 2341796-81-2 |
|  | BAY61-3606 |  | MedChemExpress | HY-14985 | 648903-57-5 |
|  | tBID | HIPK3 | MedChemExpress | HY-100464 | 1639895-85-4 |
|  | SB4614852 |  | ChemDiv | C066-2476 | 155983-19-0 |
|  | BIX02188 | MAP2K5 | MedChemExpress | HY-12055 | 334949-59-6 |
|  | BIX02189 |  | MedChemExpress | HY-12056 | 1265916-41-3 |
|  | BI8626 | HUWE1 | MedChemExpress | HY-120204 | 1875036-75-1 |
|  | BI8622 |  | MedChemExpress | HY-120929 | 1875036-74-0 |
|  | ICCB-19 | TRADD | Enamine | EN300-195751 | 750621-52-4 |
|  | TAK-243 | UBA6 | MedChemExpress | HY-100487 | 1450833-55-2 |
|  | Competitor_USP30 | USP30 | Enamine | EN300-6475230 | 2241022-48-8 |
|  | GSK8612 | TBK1 | MedChemExpress | HY-111941 | 2361659-62-1 |
| Agonists | ML-SA1 | TRPML1 | Vitas-M Laboratory, Ltd. | STK095286 | 332382-54-4 |
|  | MK6-83 |  | Sigma Aldrich | SML1509 | 1062271-24-2 |
|  | SF-22 |  | Enamine | Z45692898 | 746609-35-8 |
|  | SF-51 |  | Vitas-M Laboratory, Ltd. | STK061646 | 304869-80-5 |
|  | Cytosporone B | NR4A1 | Tocris Bioscience | 5459 | 321661-62-5 |
|  | DIM-C-pPhOCH3 |  | Tocris Bioscience | 6376 | 33985-68-1 |
|  | Celastrol |  | Key Organics | GS-6082 | 34157-83-0 |

## Author contributions

AJW, NTK and JS conceived and designed the project and acquired the funding. MLG, EC, WHA, CR, MM, O-GC, VZ and SM carried out the experimental investigation. GA assembled and curated the chemical library resource, W-LK, JHC and JD provided further conception and design and also provided supervision. AJW, NTK and MLG wrote the original draft and also reviewed and edited with further support from JS.

## Acknowledgements

The authors thank Peter Atkinson and James Allpress (representing Eisai Limited) Fiona Menzies (representing Eli Lilly and Company), Carla F. Bento, Nicola G. Wallis, Elena Di Daniel and Heather Weir (representing Astex Pharmaceuticals), and Alison Schuldt and Kathryn Chapman (representing the Milner Therapeutics Institute) for their valuable contributions to the development and progression of this project. The authors also acknowledge the financial support provided by Eisai Limited, Astex Pharmaceuticals, and Eli Lilly and Company. The ALBORADA Drug Discovery Institute acknowledges core funding from Alzheimer’s Research UK (registered charity no.1077089 and SC042474), supported by the ALBORADA Trust.

