## Supplementary Figures for "Cross-platform multi-laboratory screening identifies HUWE1, USP33 and USP20 as negative-regulators of autophagy or mitophagy with relevance to neurodegeneration"

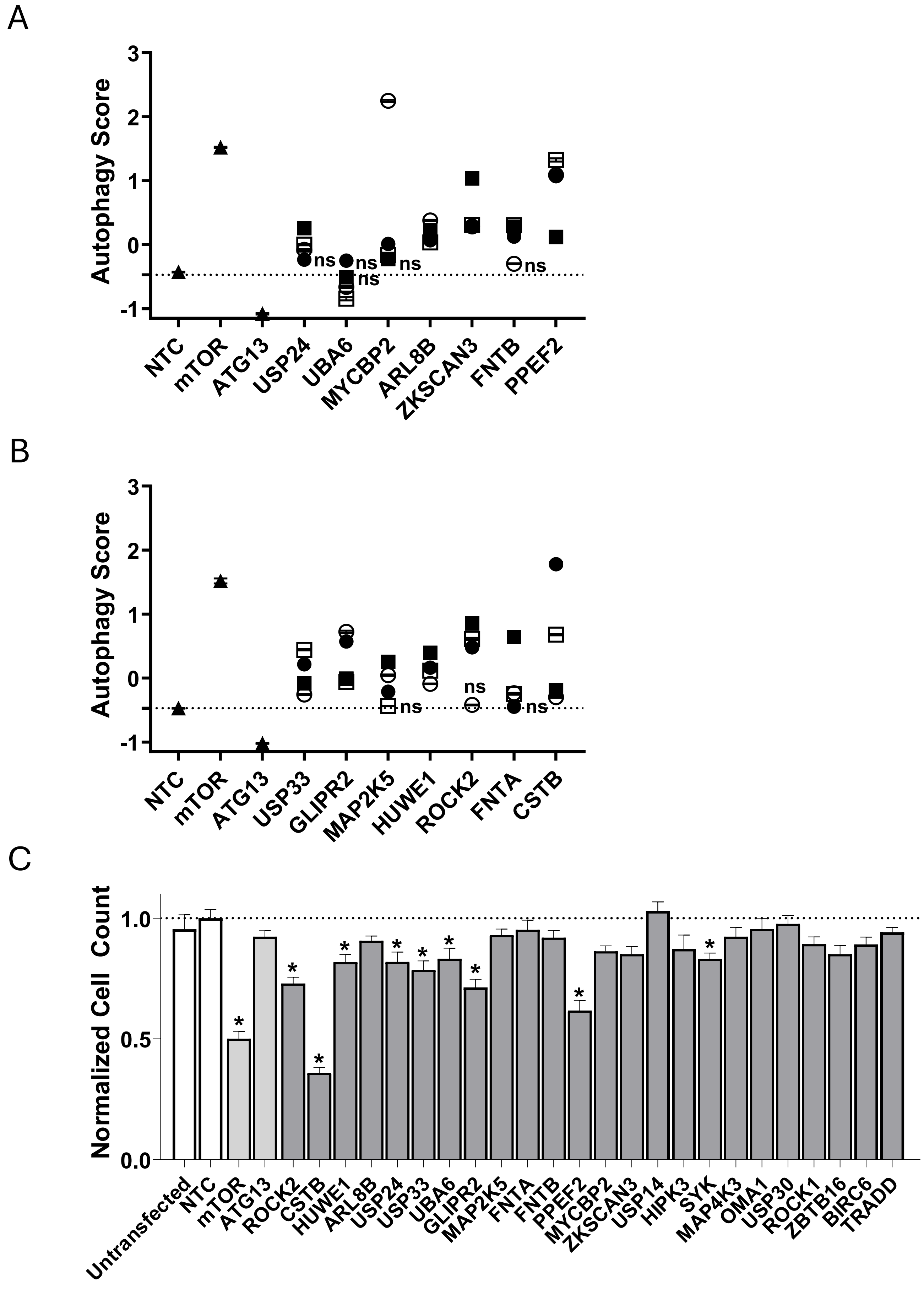
A
Autophagy Score
3
2
1
0
-1
ns
NTC
mTOR
ATG13
USP24
UBA6
MYCBP2
ARL8B
ZKSCAN3
FNTB
PPEF2
B
Autophagy Score
NTC
mTOR
ATG13
USP33
GLIPR2
MAP2K5
HUWE1
ROCK2
FNTA
CSTB
C
Normalized Cell Count
1.0
0.5
0.0
*
Untransfected
NTC
mTOR
ATG13
ROCK2
CSTB
HUWE1
ARL8B
USP24
USP33
UBA6
GLIPR2
MAP2K5
FNTA
FNTB
PPEF2
MYCBP2
ZKSCAN3
USP14
HIPK3
SYK
MAP4K3
OMA1
USP30
ROCK1
ZBTB16
BIRC6
TRADD

A

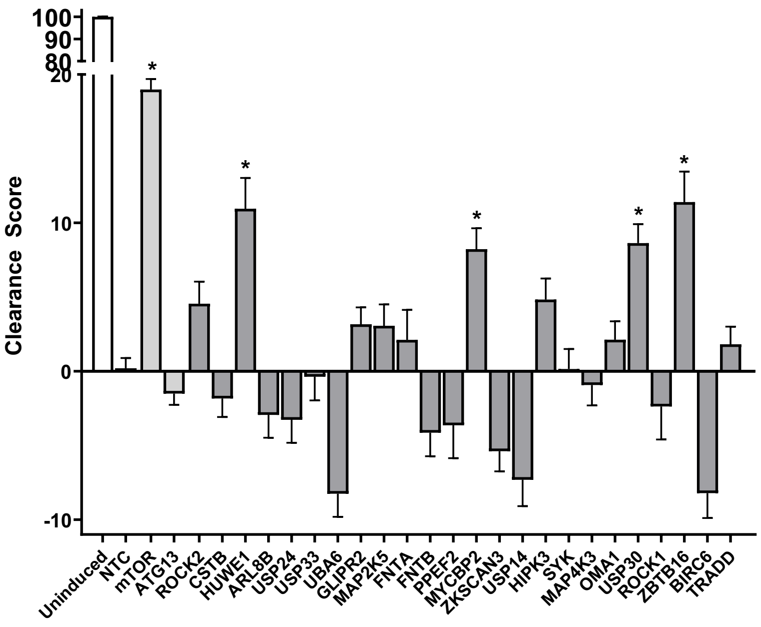
Clearance Score
100
90
80
20
10
0
-10
*
Uninduced
NTC
mTOR
ATG13
ROCK2
CSTB
HUWE1
ARL8B
USP24
USP33
UBA6
GLIPR2
MAP2K5
FNTA
FNTB
PPEF2
MYCBP2
ZKSCAN3
USP14
HIPK3
SYK
MAP4K3
OMA1
USP30
ROCK1
ZBTB16
BIRC6
TRADD

B

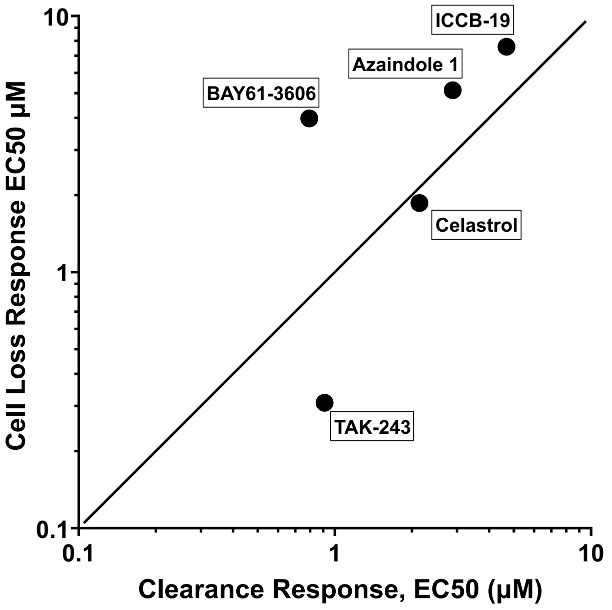
Cell Loss Response EC50 µM
Clearance Response, EC50 (µM)
ICCB-19
Azaindole 1
BAY61-3606
Celastrol
TAK-243
0.1
1
10

C

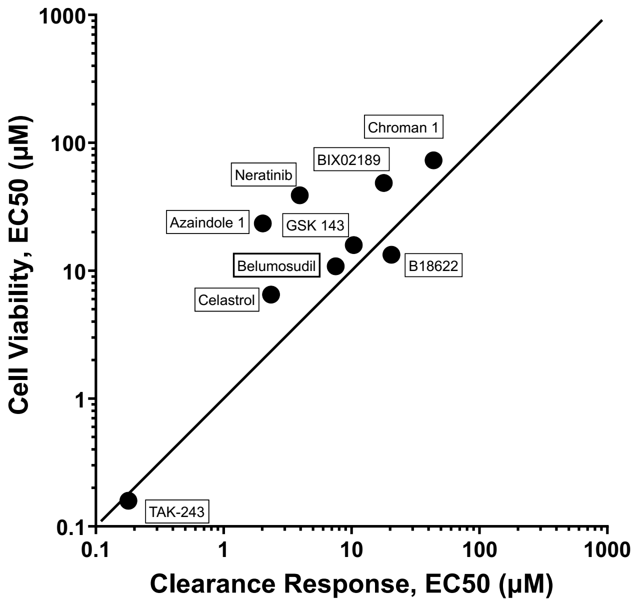
Cell Viability, EC50 (µM)
Clearance Response, EC50 (µM)
Chroman 1
BIX02189
Neratinib
Azaindole 1
GSK 143
Belumosudil
B18622
Celastrol
TAK-243
0.1
1
10
100
1000

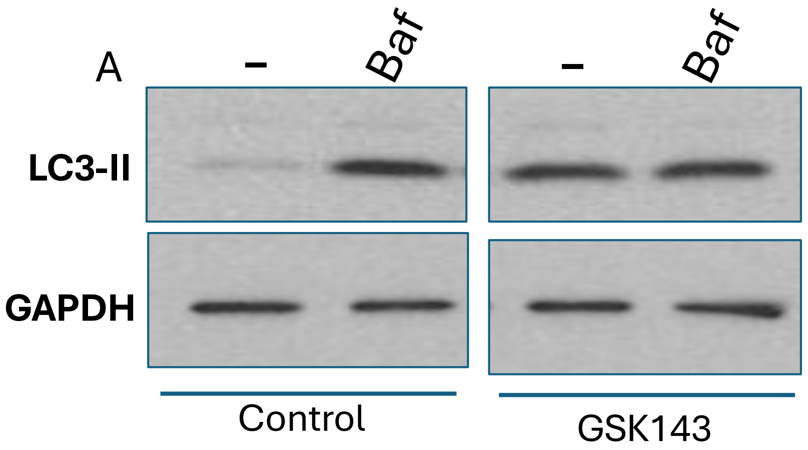
A
–
Baf
–
Baf
LC3-II
GAPDH
Control
GSK143

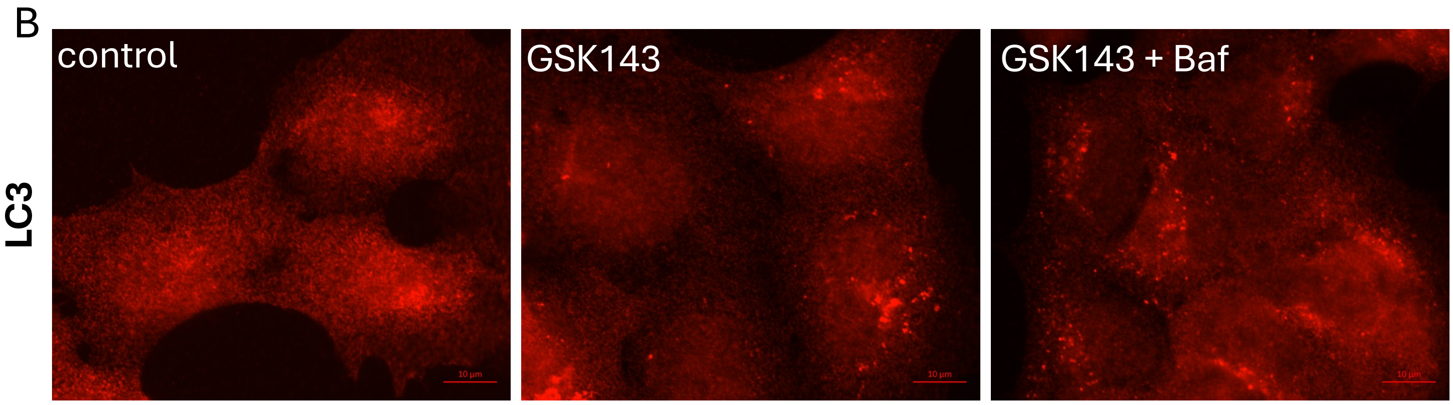
B
LC3
control
GSK143
GSK143 + Baf

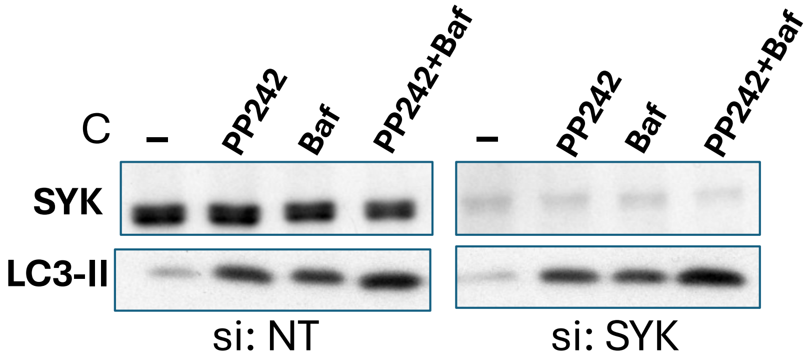
C
–
PP242
Baf
PP242+Baf
–
PP242
Baf
PP242+Baf
SYK
LC3-II
si: NT
si: SYK

MAP2K5 SUMMARY DATA

Suppl Figure 4

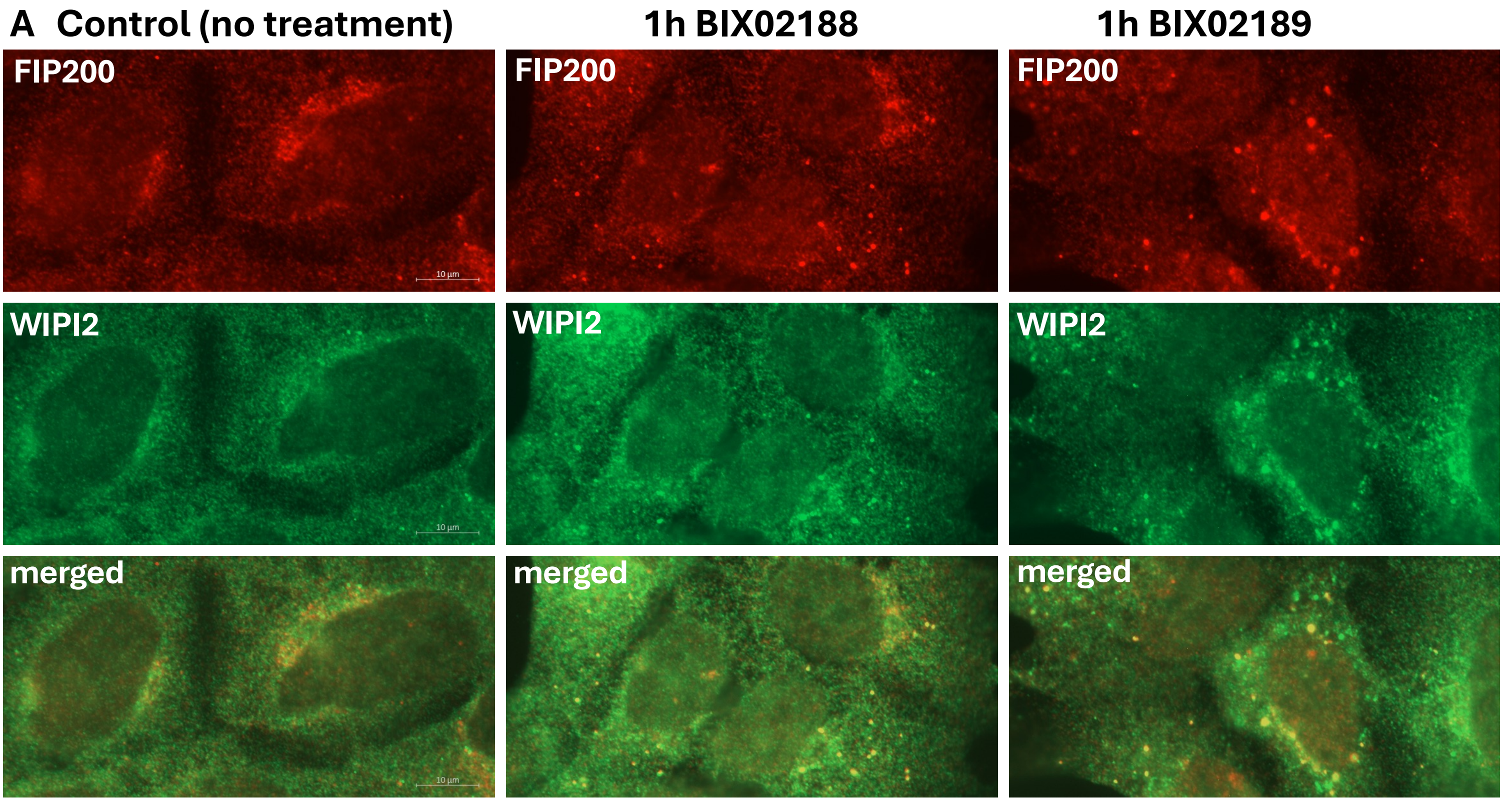


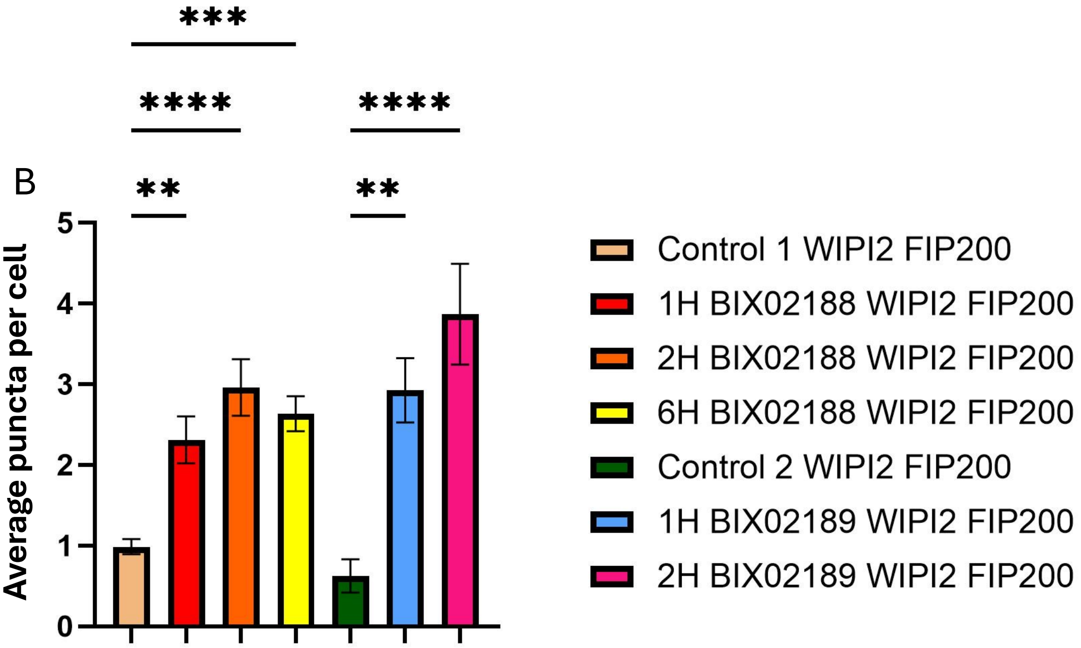


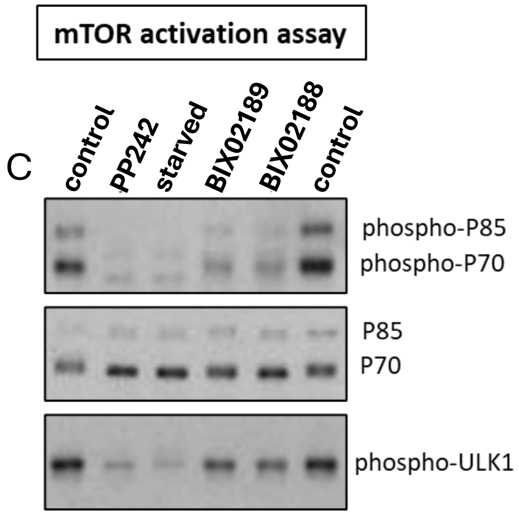

### ARL8B SUMMARY DATA

Suppl Figure 5

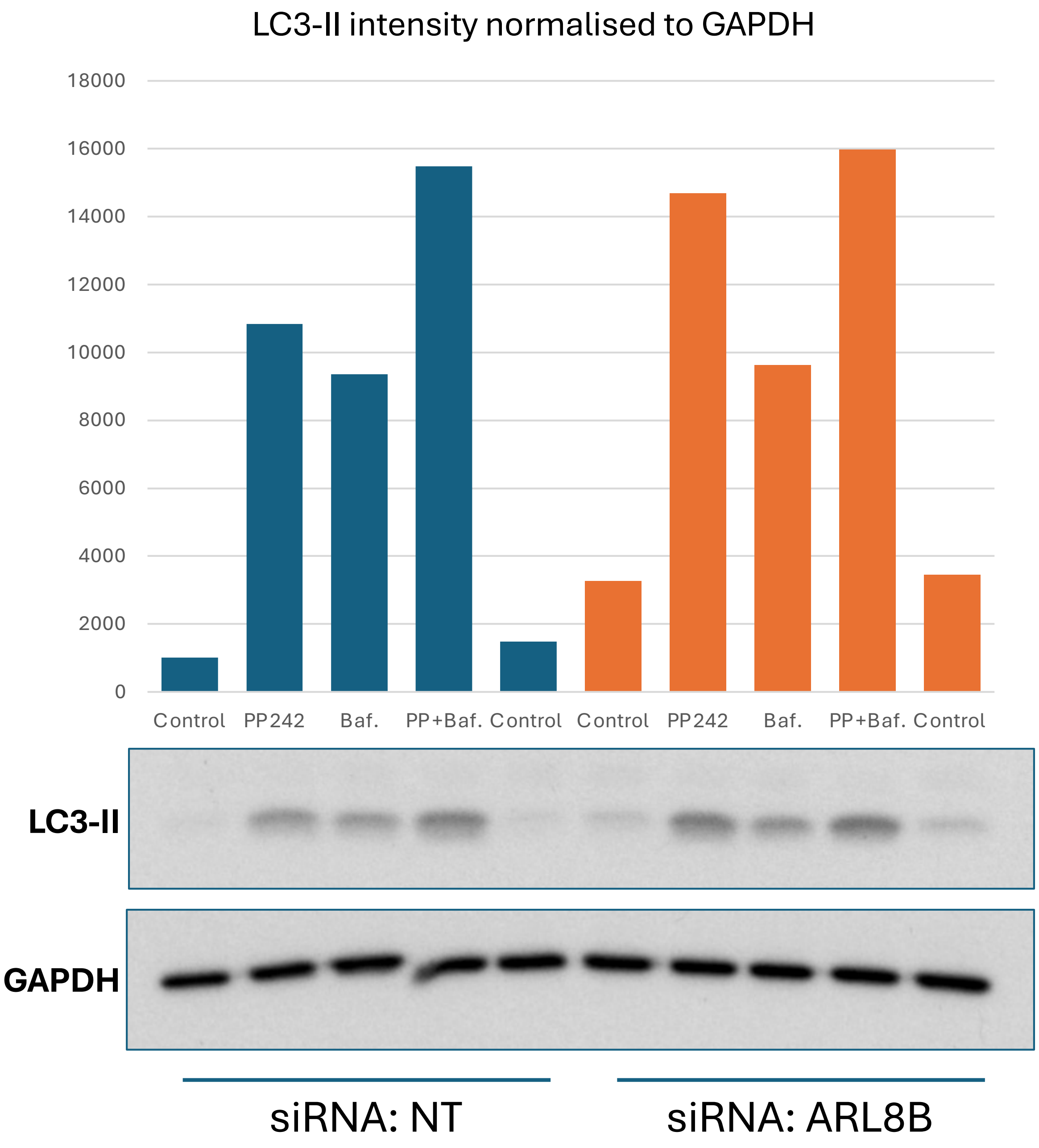

### TRPML SUMMARY DATA

Suppl Figure 6

A

Number of LC3 puncta per cell with various TRPML compounds.

| | 1 hr | | 15 hr | |
|---|---|---|---|---|
| | WT | ATG13-KO | WT | ATG13-KO |
| DMSO | 0 | 0 | 0 | 0 |
| PP242 | 3 | 0 | 3 | 0 |
| SF-22 | 3 | 3 | 3 | 2 |
| SF-51 | 2 | 2 | 3 | 1 |
| MK6-83 | 3 | 3 | 3 | 3 |
| ML-SA1 | 3 | 3 | 3 | 3 |

B

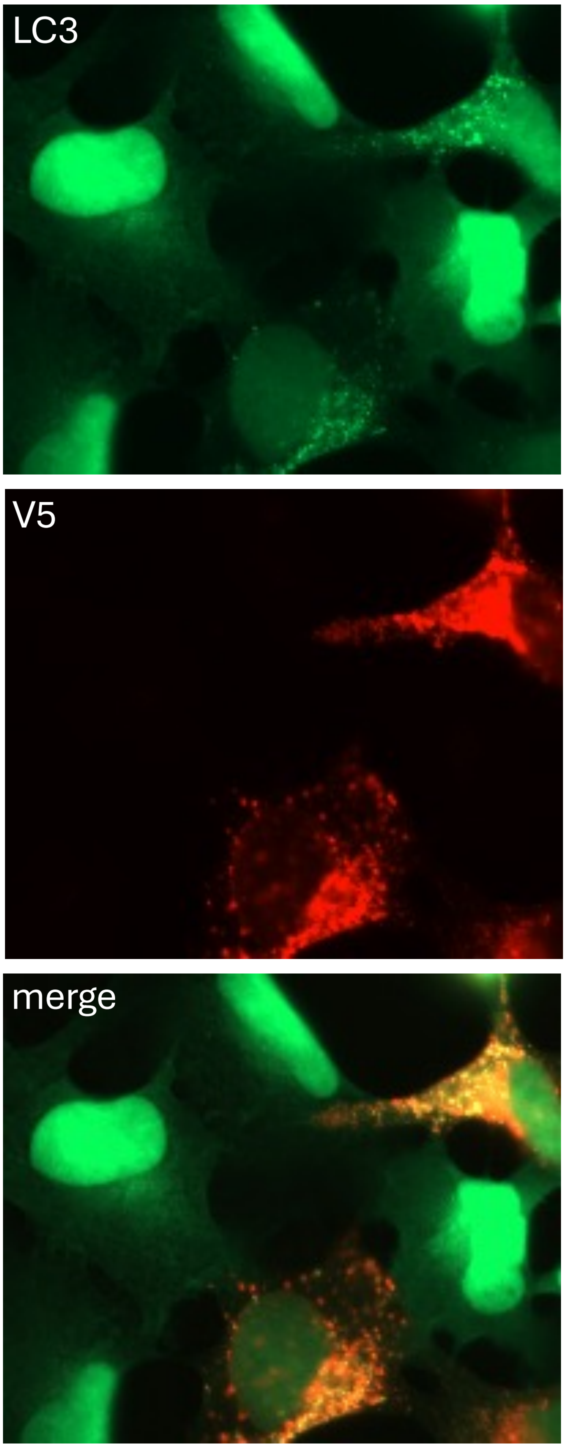


C

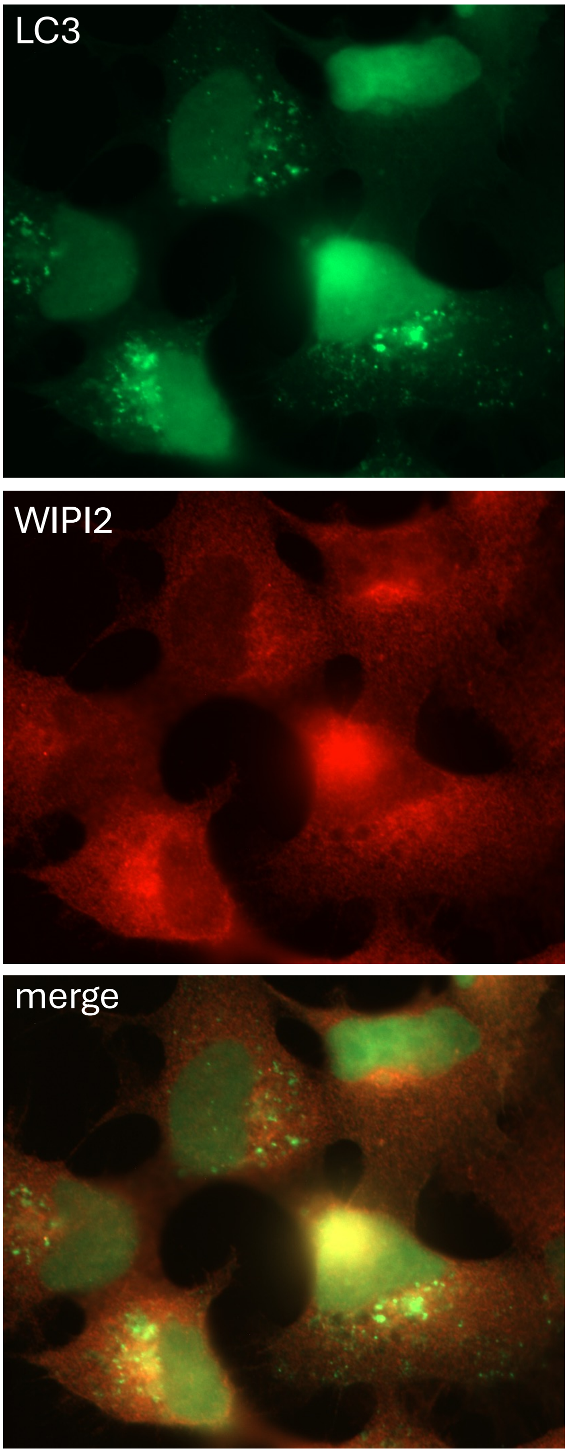

USP24 SUMMARY DATA

Suppl Figure 7

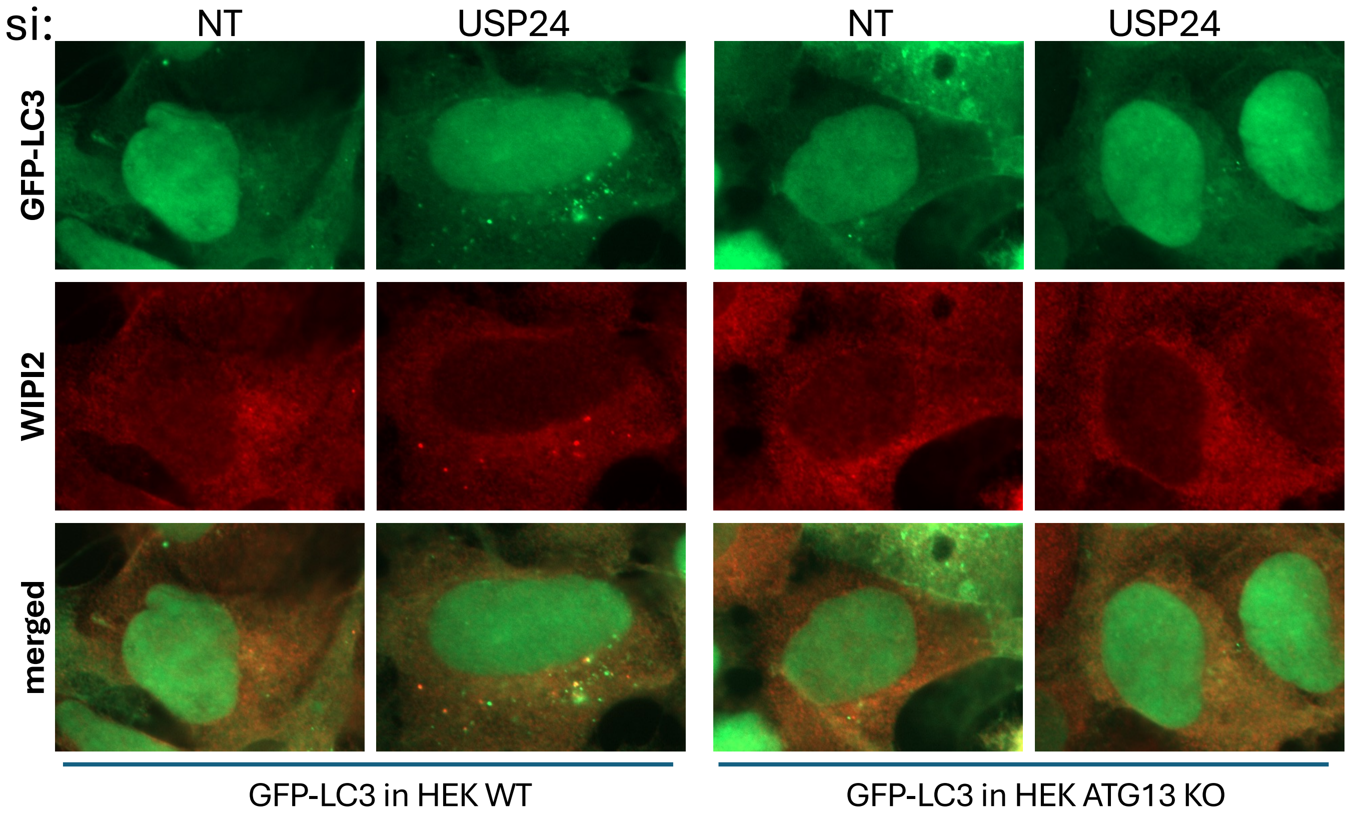

ROCK2 SUMMARY DATA



A

ROCK2 siRNA in GFP-LC3 HEK cells

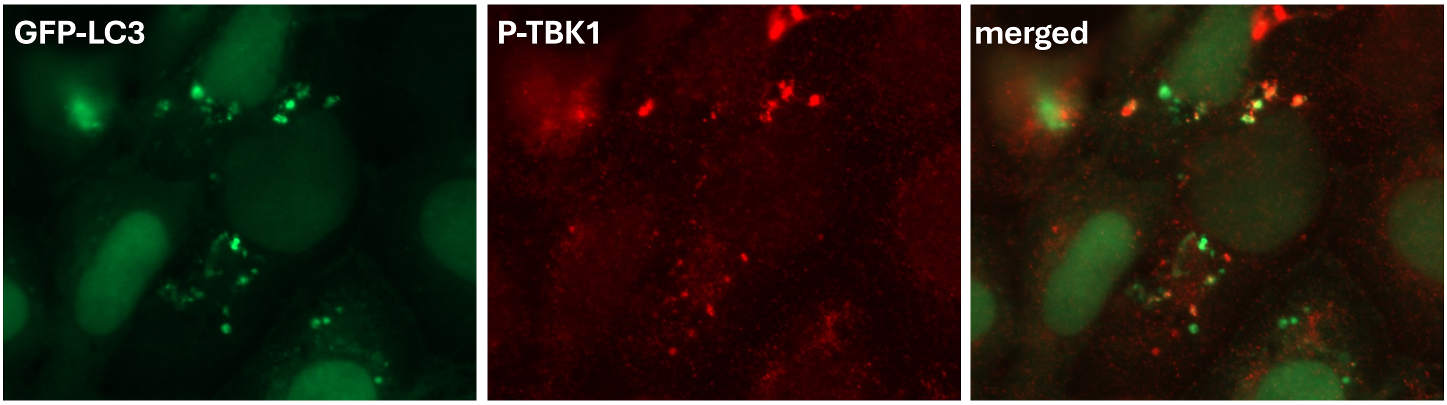


B

ROCK2 siRNA in GFP-ATG13 HEK cells

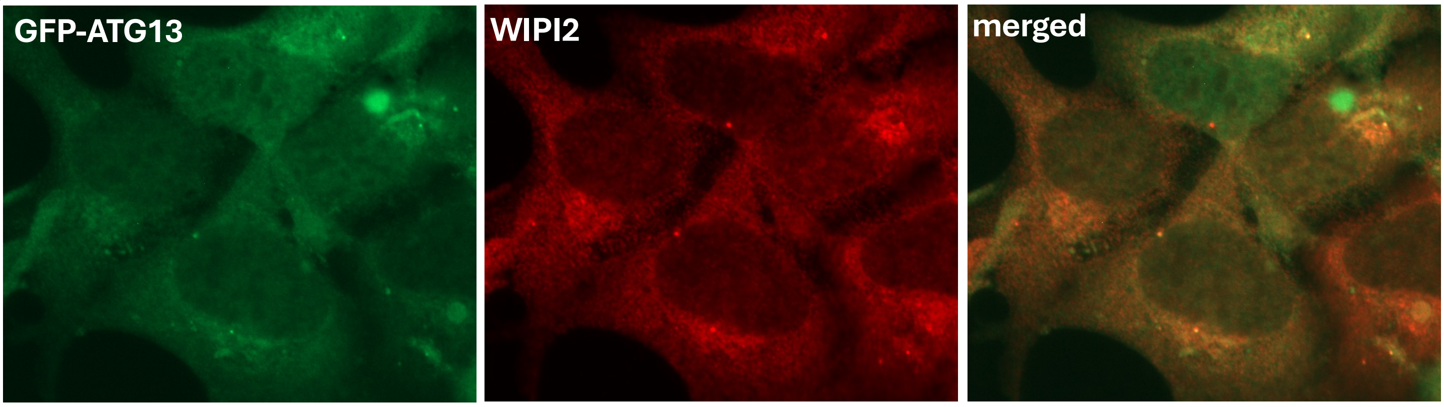


C

GFP-LC3 HEK cells treated with ROCK2 inhibitor Belumosudil

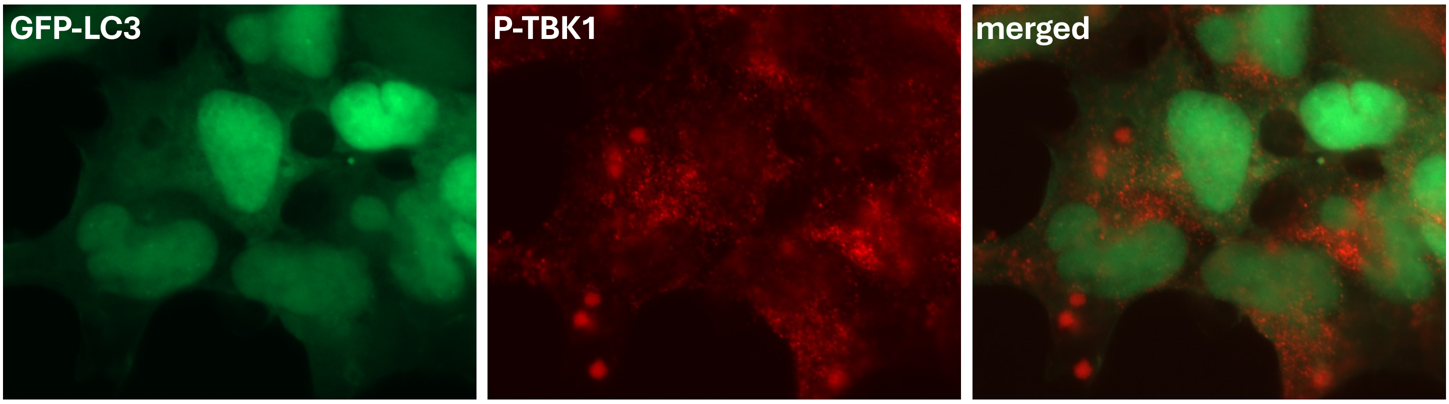

USP 20/33 SUMMARY DATA

Suppl Figure 9

A

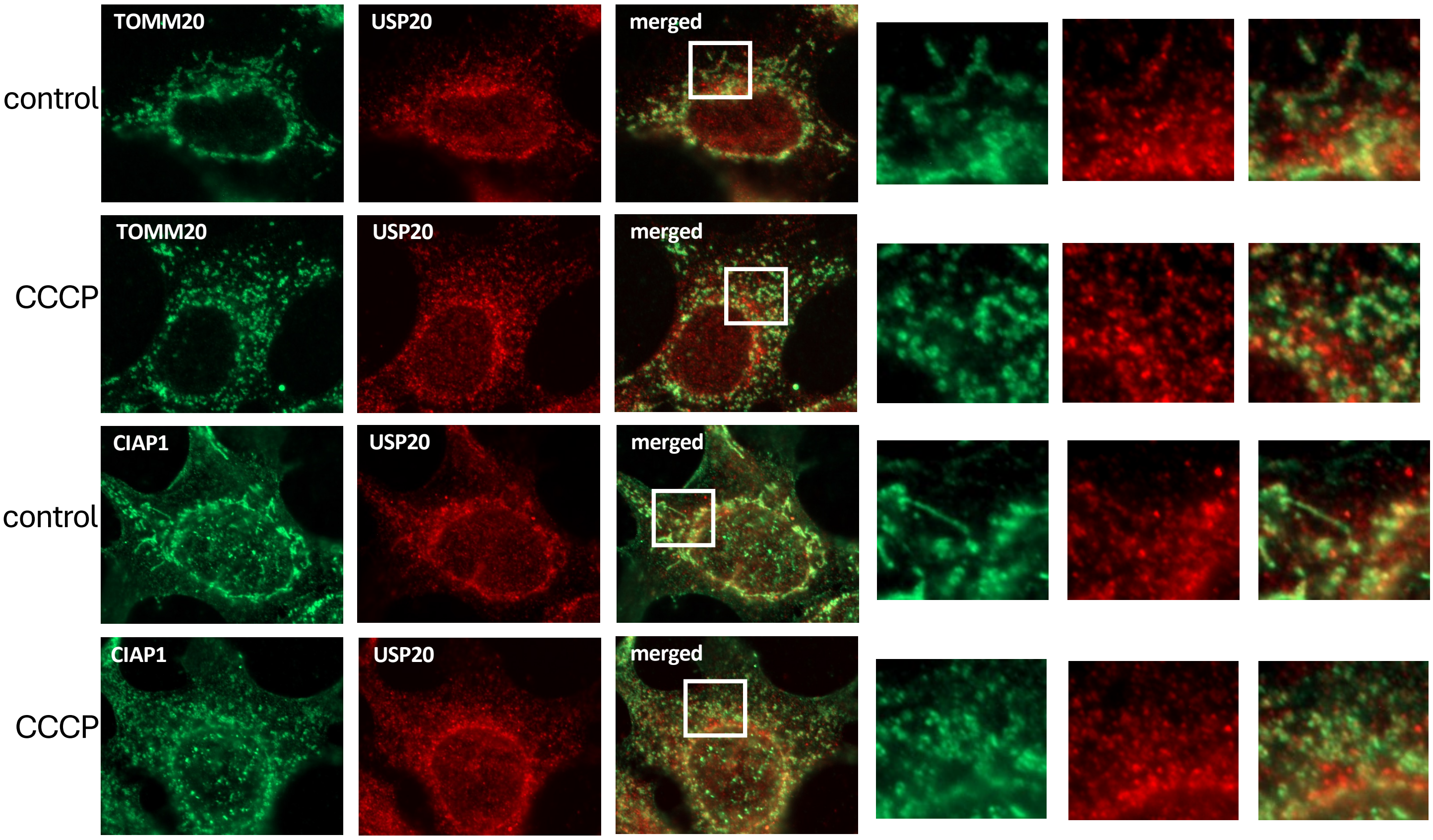


B

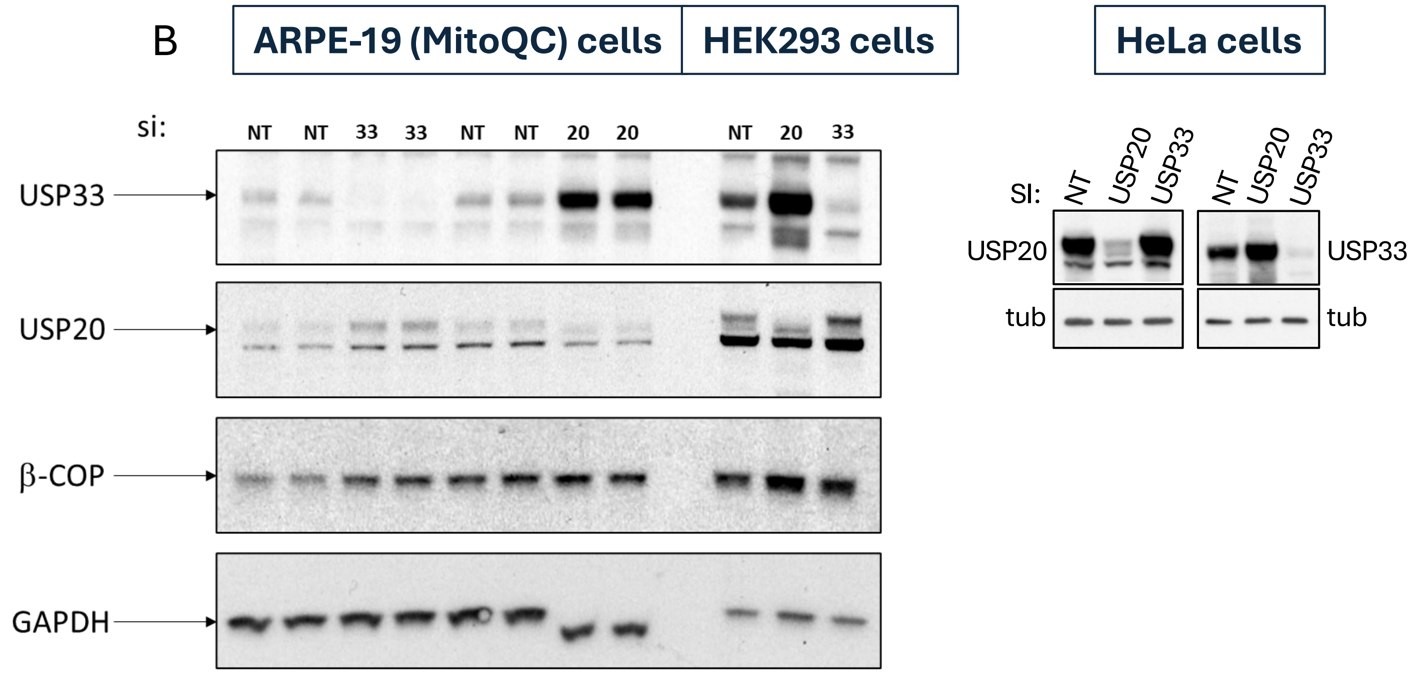

USP30 SUMMARY DATA



HEK-293 cells expressing GFP-LC3

A

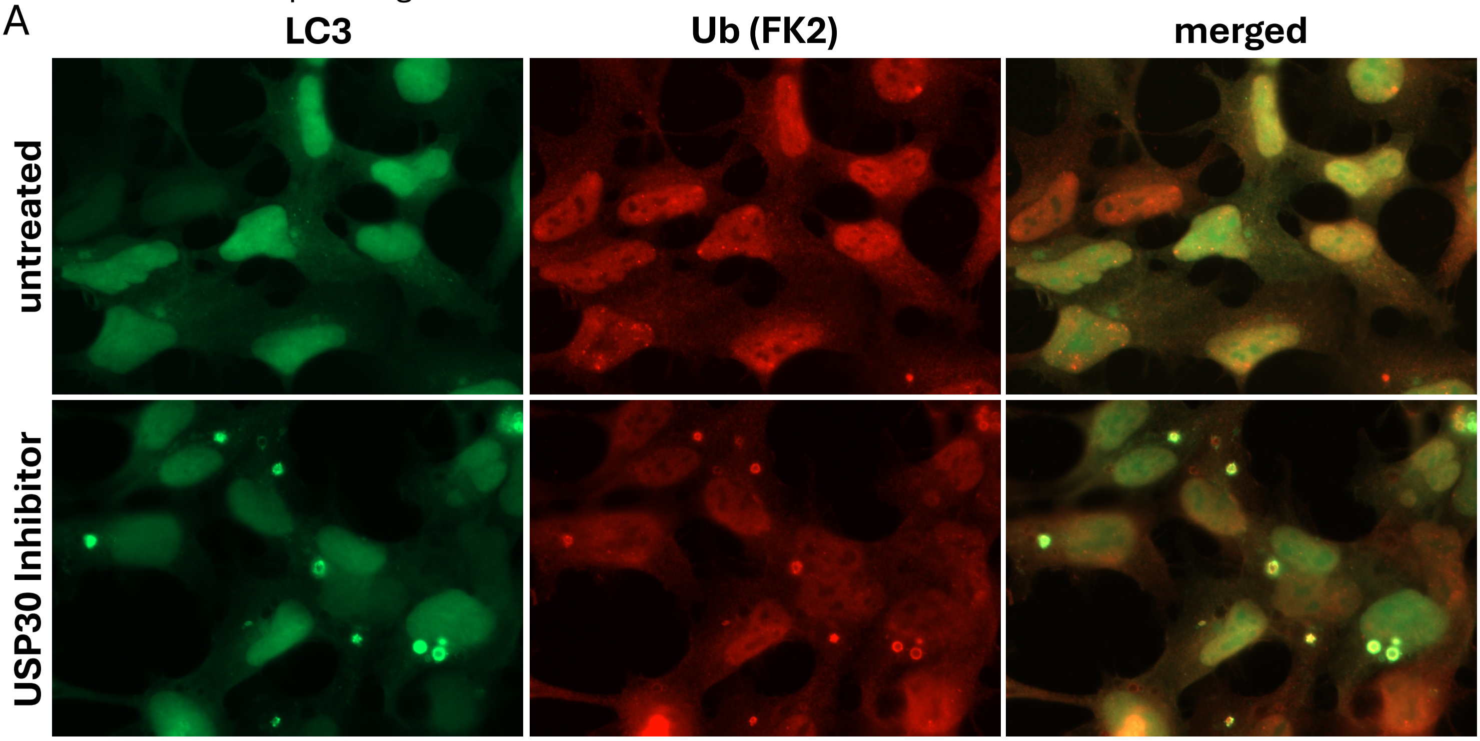


HEK-293 cells expressing GFP-ATG13

B

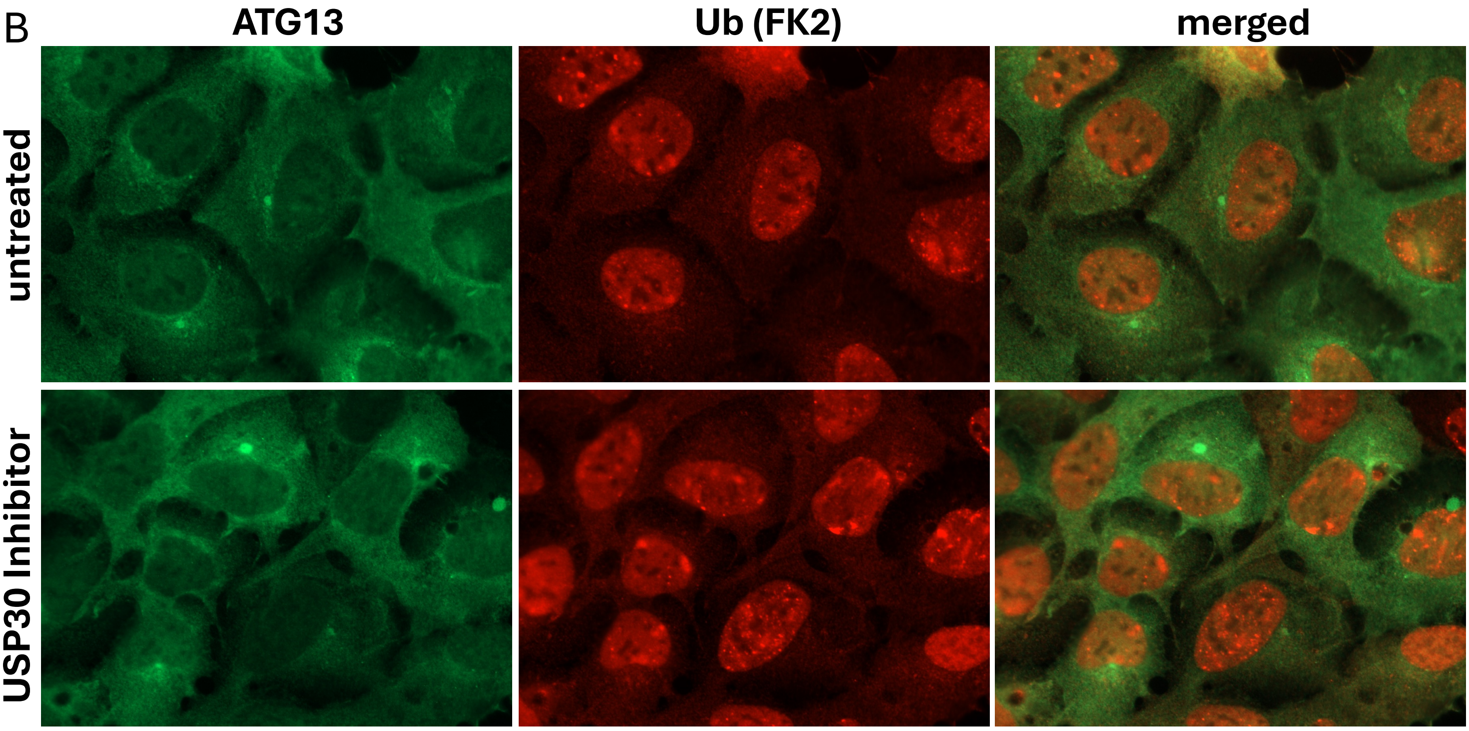


HEK-293 cells expressing GFP-LC3

C

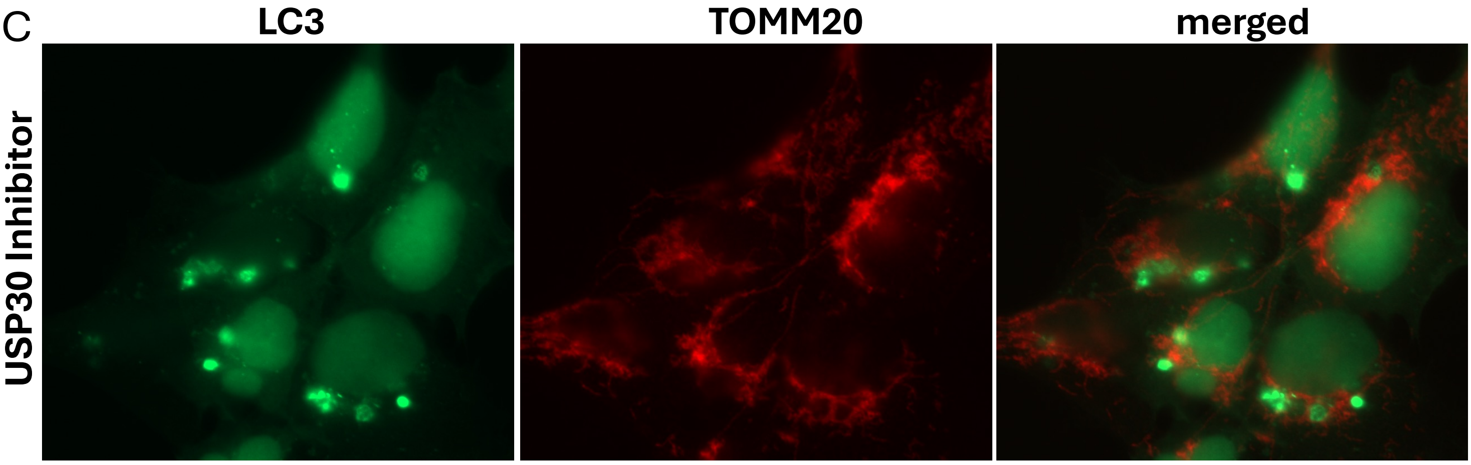
